# Border-zone cardiomyocytes and macrophages contribute to remodeling of the extracellular matrix to promote cardiomyocyte invasion during zebrafish cardiac regeneration

**DOI:** 10.1101/2024.03.12.584570

**Authors:** Florian Constanty, Bailin Wu, Ke-Hsuan Wei, I-Ting Lin, Julia Dallmann, Stefan Guenther, Till Lautenschlaeger, Rashmi Priya, Shih-Lei Lai, Didier Y.R. Stainier, Arica Beisaw

## Abstract

Despite numerous advances in our understanding of zebrafish cardiac regeneration, an aspect that remains less studied is how regenerating cardiomyocytes invade, and eventually replace, the collagen-containing fibrotic tissue following injury. Here, we provide an in-depth analysis of the process of cardiomyocyte invasion using live-imaging and histological approaches. We observed close interactions between protruding cardiomyocytes and macrophages at the wound border zone, and macrophage-deficient *irf8* mutant zebrafish exhibited defects in extracellular matrix (ECM) remodeling and cardiomyocyte protrusion into the injured area. Using a resident macrophage ablation model, we show that defects in ECM remodeling at the border zone and subsequent cardiomyocyte protrusion can be partly attributed to a population of resident macrophages. Single-cell RNA-sequencing analysis of cells at the wound border revealed a population of cardiomyocytes and macrophages with fibroblast-like gene expression signatures, including the expression of genes encoding ECM structural proteins and ECM-remodeling proteins. The expression of *mmp14b*, which encodes a membrane-anchored matrix metalloproteinase, was restricted to cells in the border zone, including cardiomyocytes, macrophages, fibroblasts, and endocardial/endothelial cells. Genetic deletion of *mmp14b* led to a decrease in 1) macrophage recruitment to the border zone, 2) collagen degradation at the border zone, and 3) subsequent cardiomyocyte invasion. Furthermore, cardiomyocyte-specific overexpression of *mmp14b* was sufficient to enhance cardiomyocyte invasion into the injured tissue and along the apical surface of the wound. Altogether, our data shed important insights into the process of cardiomyocyte invasion of the collagen-containing injured tissue during cardiac regeneration. They further suggest that cardiomyocytes and resident macrophages contribute to ECM remodeling at the border zone to promote cardiomyocyte replenishment of the fibrotic injured tissue.

## Introduction

Despite numerous advances in our understanding of cardiovascular biology, heart disease remains the leading cause of morbidity and mortality worldwide^1, 2^. Following acute myocardial infarction in the human heart, millions of cardiomyocytes are lost and replaced with a fibrotic scar. While this fibrotic scar is necessary to maintain the structural integrity of the heart, its presence, in combination with decreased contractile mass, eventually leads to heart failure. There remains a pressing need to develop therapeutic strategies to replace lost heart tissue. Non-mammalian vertebrate organisms, such as the teleost zebrafish, have the ability to regenerate lost cardiomyocytes following multiple types of injury^3, 4^. Notably, zebrafish can regenerate their hearts following cryoinjury, an injury model whereby a liquid nitrogen-cooled probe placed on the heart leads to cell death, activation of an inflammatory response, and deposition of a fibrotic collagen-containing scar, similar to myocardial infarction in the human heart^5–7^. While research efforts in the last two decades have led to an immense increase in our knowledge of the mechanisms of cardiac regeneration^8^, it remains unclear how regenerating cardiomyocytes at the wound border invade and eventually replace the collagen-containing scar tissue following injury.

Previous studies in zebrafish using photo-convertible Kaede protein have reported that zebrafish cardiomyocytes on the surface of the heart migrate, or are displaced, from the wound border to the injured area during the regeneration process^9^. This migration was shown to rely on the chemokine receptor Cxcr4, as treatment of adult fish with a CXCR4 antagonist following cardiac injury abrogated the presence of photo-converted cardiomyocyte-specific Kaede in the injured area. Furthermore, treatment with a CXCR4 antagonist led to a defect in scar resolution without affecting cardiomyocyte proliferation^9^. In neonatal mouse hearts, which can also regenerate following resection of the apex, cardiomyocytes protruding into the wound were shown to be essential for cardiac regeneration^10^. These protrusions were shown to rely on the dystrophin glycoprotein complex that links the actin cytoskeleton to the extracellular matrix (ECM), as mutant cardiomyocytes harboring a null *Dystrophin* mutation (from *mdx* mutant mice) lacked cardiomyocyte protrusions and isolated *mdx* mutant cardiomyocytes were not able to migrate into a collagen gel. Despite normal cardiomyocyte proliferation, *mdx* mutant mice retained a fibrotic scar^10^. These studies indicate that cardiomyocyte migration and invasion are necessary for resolution of the fibrotic scar during the cardiac regeneration process. Furthermore, results of these published studies suggest that cardiomyocyte proliferation alone is not sufficient to promote scar resolution following injury.

Notably, in two genetic mouse models that can promote regeneration in the nonregenerative adult mouse heart, cardiomyocyte protrusion and migration were also observed: first, activation of the transcription factor YAP in cardiomyocytes from YAP5SA overactivation or *Hippo*-deficient genetic models leads to cardiomyocyte proliferation and regeneration in the adult mouse heart^11–13^. YAP binds to several target genes to activate their expression in cardiomyocytes following myocardial infarction, including those that encode regulators of cytoskeletal dynamics^10^. *Hippo*-deficient cardiomyocytes extend protrusions into the infarcted area and isolated *Hippo-*deficient cardiomyocytes are able to migrate through a collagen gel matrix *in vitro*^10^. In another genetic model, cardiomyocyte-specific expression of a constitutively active form of *Erbb2* (caERBB2) is able to promote cardiomyocyte regeneration in the adult mouse heart^14^. Mechanistic analyses revealed that caERBB2 drives the activation of YAP, promotes the migration of isolated cardiomyocytes *in vitro*, and upregulates the expression of several genes that encode regulators of the extracellular matrix, including *Loxl2, Mmp14, Mmp2,* and *Pcolce*^15^. These studies suggest that cardiomyocyte protrusion and migration are regulated by genetic programs that can promote cardiac regeneration in the adult mammalian heart.

Following cryoinjury in the zebrafish heart, multiple cell types orchestrate the response to injury^8^. For example, endothelial, endocardial, and epicardial cells have been shown to provide growth factors and signaling molecules to promote revascularization of the injured tissue and cardiomyocyte regeneration^16–21^. Fibroblasts, which largely arise from the epicardium, have been shown to deposit ECM to support the heart during the process of cardiac regeneration, and are essential for cardiomyocyte proliferation following injury^22–25^. The importance of immune cells in promoting the cardiac regeneration process has also been recognized in more recent years. In particular, macrophages have been shown to play an essential role in regeneration of zebrafish, salamander, and neonatal mouse hearts in response to injury^26–29^. Macrophages have been shown to directly contribute to and regulate the composition of scar/ECM in the cryoinjured heart^27, 30, 31^. Furthermore, depletion of macrophages by clodronate liposome administration or the use of genetic models leads to defects in CM proliferation and neovascularization in regenerating hearts^26, 28, 32, 33^. While it is clear that many, if not all, cardiac cell types play an essential role in the process of cardiac regeneration, their contribution to cardiomyocyte repopulation of fibrotic tissue following cryoinjury is largely unknown.

Here, we provide an in-depth analysis of cardiomyocyte protrusion into the injured area during zebrafish cardiac regeneration. We show that cardiomyocytes at the wound border exhibit characteristics of migratory cells, including motile filopodia-like extensions into the injured tissue and upregulation of gene expression programs regulating actin dynamics, focal adhesions, and ECM remodeling. Furthermore, we show that macrophages play an essential role at the border zone to remodel the ECM and promote cardiomyocyte invasion. Lastly, we show that Mmp14b is an important regulator of macrophage presence and ECM remodeling at the border zone, and subsequent CM invasion into injured tissue during cardiac regeneration in zebrafish.

## Results

### Characterization of cardiomyocyte protrusion and invasion into the injured tissue during zebrafish cardiac regeneration

While previous studies have indicated that cardiomyocyte (CM) protrusion and invasion into the injured area are likely essential for cardiac regeneration^9, 10, 15^, we lack a detailed knowledge of the molecular mechanisms underlying this process. To characterize CM protrusion and invasion, we performed a time-course analysis at multiple timepoints following cryoinjury. Using phalloidin, which labels actin filaments (F-actin) and is highly abundant in zebrafish cardiomyocytes (**Supp. Fig. 1A and 1B**), we observed a peak in the number of CM protrusions at the wound border zone between 7 and 10 days post cryoinjury (dpci, **Fig. 1A** and **1B**). Furthermore, we observed a peak in CM protrusion length at 7 dpci (**Fig. 1B**), corresponding to timepoints post cryoinjury that exhibit high levels of CM proliferation. To determine the relationship between protruding and proliferating CMs, we utilized the FUCCI (fluorescent ubiquitination-based cell cycle indicator) cell-cycle reporter line *Tg(myl7:mVenus-gmnn); Tg(myl7:mCherry-cdt1)*, marking CMs in S/G2/M phase with mVenus and CMs in G0/G1 phase with mCherry. Immunostaining of ventricular sections at 10 dpci revealed that approximately 70% of CM nuclei adjacent to CM protrusions into the injured area are in G0/G1 phase (**Fig. 1C**). These observations indicate that CM protrusion and proliferation are regenerative processes that are spatially uncoupled, and suggest that CM proliferation may precede protrusion into the injured tissue. Additional processes that have been described during cardiac regeneration include cardiomyocyte dedifferentiation and sarcomere disassembly^34–37^. We examined *Tg(gata4:EGFP)* ventricular sections at 10 dpci and found that nearly all CMs protruding into the injured area exhibited *gata4*:EGFP localization, indicating that CMs that invade the injured area have undergone dedifferentiation (**Fig. 1D**). Further, we examined CM sarcomere structure using ventricular sections from *Tg(myl7:actn3b-EGFP)* zebrafish at 10 dpci and observed sarcomere disorganization and disassembly at the distal tips of CMs protruding into the injured area (**Fig. 1E**), in line with our previously published data^38^. Lastly, we examined ventricular sections from *Tg(myl7:LIFEACT-GFP)* zebrafish at 10 dpci and found that invading CMs at the border zone extend actin-filled protrusions into the injured area (**Fig. 1F**).

**Figure 1:**
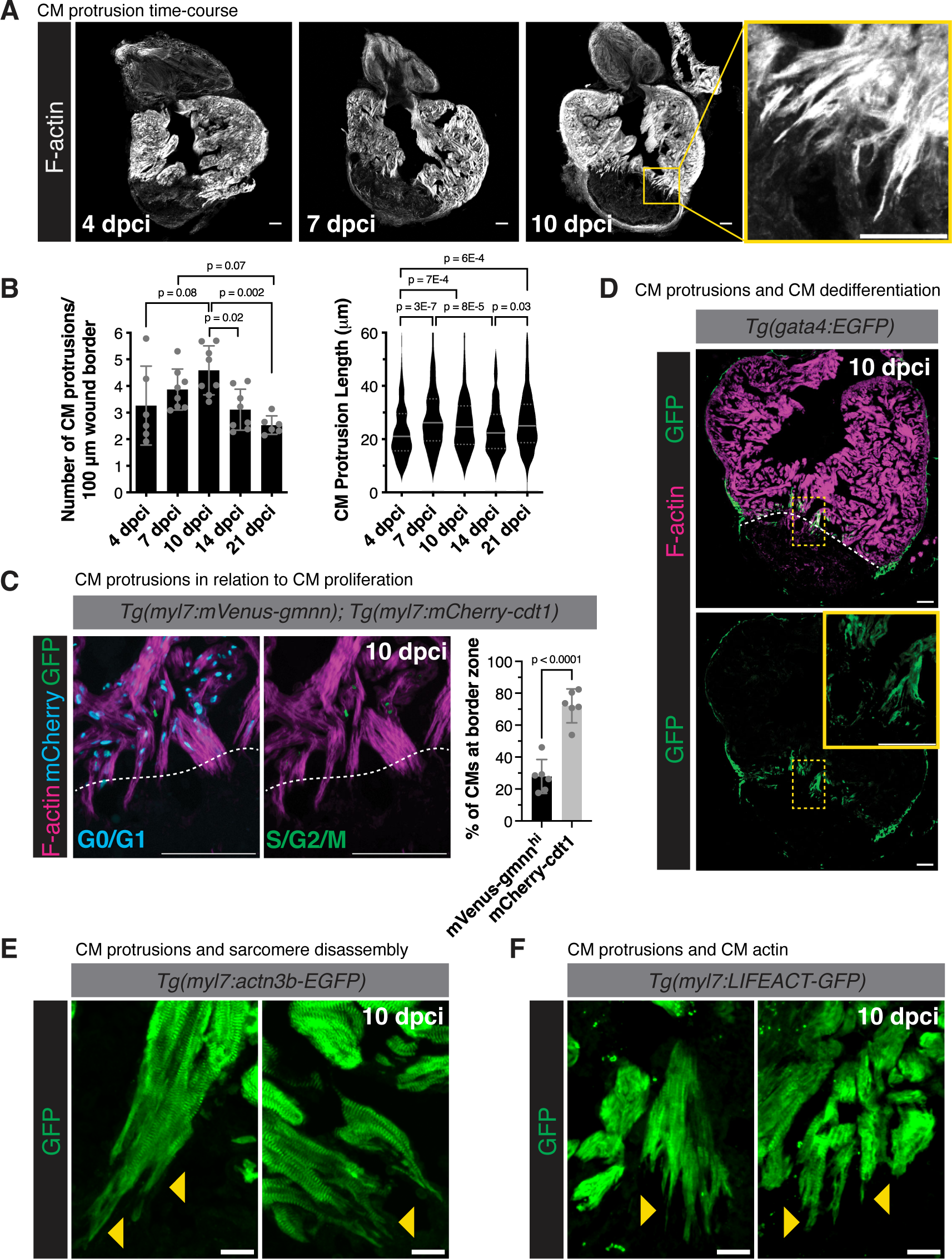
Characterization of CM protrusion into the injured area during zebrafish heart regeneration. **(A)** Phalloidin staining of F-actin in thick cryosections of zebrafish ventricles at 4, 7, and 10 days post cryoinjury (dpci). Yellow box denotes the zoomed image from the wound border zone at 10 dpci. **(B)** Quantification of the number of CM protrusions per 100 micron of wound border (left) and length of CM protrusions (right) from thick cryosections of adult ventricles at 4 (n=245 CMs from 6 ventricles), 7 (n=323 CMs from 8 ventricles), 10 (n=449 CMs from 8 ventricles), 14 (n=300 CMs from 8 ventricles), and 21 (n=248 CMs from 6 ventricles) dpci. P-values were calculated using one-way ANOVA with Tukey’s multiple comparisons test (number of CM protrusions) and the Kruskal-Wallis test with Dunnett’s multiple comparisons test (CM protrusion length). **(C)** Immunostaining of GFP, mCherry, and F-actin in *Tg(myl7:mVenus-gmnn); Tg(myl7:mCherry-cdt1)* ventricles at 10 dpci. White dashed line indicates the injury border. The percentages of mVenus-Gmnn^hi^ and mCherry-Cdt1+ CM nuclei directly neighboring the wound border were quantified on the right in n=7 ventricles. P-value was calculated using an unpaired t-test. **(D)** GFP and Phalloidin staining of *Tg(gata4:EGFP)* ventricles at 10 dpci. White dashed line indicates the injury border and the yellow box denotes the zoomed image from the wound border zone.**(E)** GFP staining in *Tg(myl7:actn3b-EGFP)* ventricles at 10 dpci marking the CM sarcomere. Yellow arrowheads point to CM protrusions devoid of organized sarcomere structures. **(F)** GFP staining of *Tg(myl7:LIFEACT-GFP)* ventricles at 10 dpci marking CM-specific actin. Yellow arrowheads point to actin-filled CM protrusions. Scale bars: 100 μm in **(A)**, **(C)**, and **(D)**, 20 μm in **(E)** and **(F)**.

To visualize the process of CM invasion *in situ*, we optimized a previously published protocol to image living ventricular slices from regenerating zebrafish hearts (**Fig. 2A**)^37^. First, we observed that *Tg(myl7:*LIFEACT-GFP) levels were particularly high in cortically-located CMs at the border zone (**Fig. 2B**). We subjected ventricular slices from *Tg(myl7:LIFEACT-GFP)* zebrafish at 10 dpci to live imaging and found that cortically-located CMs extended/retracted filopodia-like structures into and away from the injured collagenous tissue (**Fig. 2C** and **Supp. Video 1**). We manually tracked the ends of CM protrusions at the border zone over the entire 12-hour time-lapse imaging period and found that the mean of cortically-located CM protrusion displacement was 6.447 μm, with a maximum displacement of 26.17 μm (**Fig. 2D**, Supp. Fig. 2C and **Supp. Video 2**). The majority of this displacement (65%) was in a net (-) direction, or away from the injured area (compared to 22% of displacement in a net (+) direction, or into the injured area, **Fig. 2E**). Movement of CM protrusions in a direction away from the injured tissue may reflect the previously described role for filopodia in environment sensing^39^; however, the 22% of CM protrusion movement into the injured tissue appears sufficient to promote CM invasion, as we observe that cortically-located CMs exhibit high levels of invasion into the injured tissue in fixed ventricular sections at 7 and 10 dpci (**Fig. 1D** and **Fig. 2B**). In order to follow trabecular CM protrusion over time, we used living ventricular slices from *Tg(myl7:actn3b-EGFP)* zebrafish at 10 dpci, as we found that CM Actn3b-EGFP levels were higher in trabecular CMs at the border zone (**Fig. 2B**). Trabecular CMs at 10 dpci extended/retracted filopodia-like structures into and away from the injured collagenous tissue similar to cortically-located CMs (**Supp. Fig. 2A** and **Supp. Video 3**), but with a mean of protrusion displacement of 2.418 μm, and a maximum displacement of 9.063 μm (**Fig. 2D**). However, the majority of trabecular CM protrusion displacement was in a net (+) direction (46% compared to 30% in a net (-) direction) (**Fig. 2E**). We further tracked trabecular CM protrusion at 3 dpci (**Supp. Fig. 2B** and **Supp. Video 4**) and found that the mean of protrusion displacement was lower than that of 10 dpci (1.641 μm) and that 41% of protrusions had a net displacement less than 1 μm (net (0)) (**Fig. 2E**). Notably, the total distance traveled by cortically-located and trabecular CM protrusions did not correlate with track displacement, i.e., trabecular CMs at 10 dpci traveled the most distance over the measured time-course (mean = 49.89 μm, compared to mean = 40.82 μm in cortically-located CM protrusions and mean = 41.15 μm in trabecular CM protrusions at 3 dpci) despite having less track displacement (**Fig. 2F**). In line with these measurements, the confinement ratio (net distance/total distance traveled) was highest in cortically-located CM protrusions, suggesting that cortical CM protrusions were most efficient in being displaced from their initial location (**Supp. Fig. 2D**). Lastly, we measured the maximum speed of CM protrusion movement and found that cortically-located CM protrusions moved at a maximum rate of 0.26 μm/min and trabecular CM protrusions with a maximum rate of 0.22 μm/min (**Fig. 2G**). Trabecular CM protrusions at 3 dpci moved more slowly, at a maximum rate of 0.18 μm/min (**Fig. 2G**). Altogether, these observations suggest that CMs at the border zone extend motile protrusions directed into the injured area that are reminiscent of filopodia on the leading edge of migrating cells.

**Figure 2:**
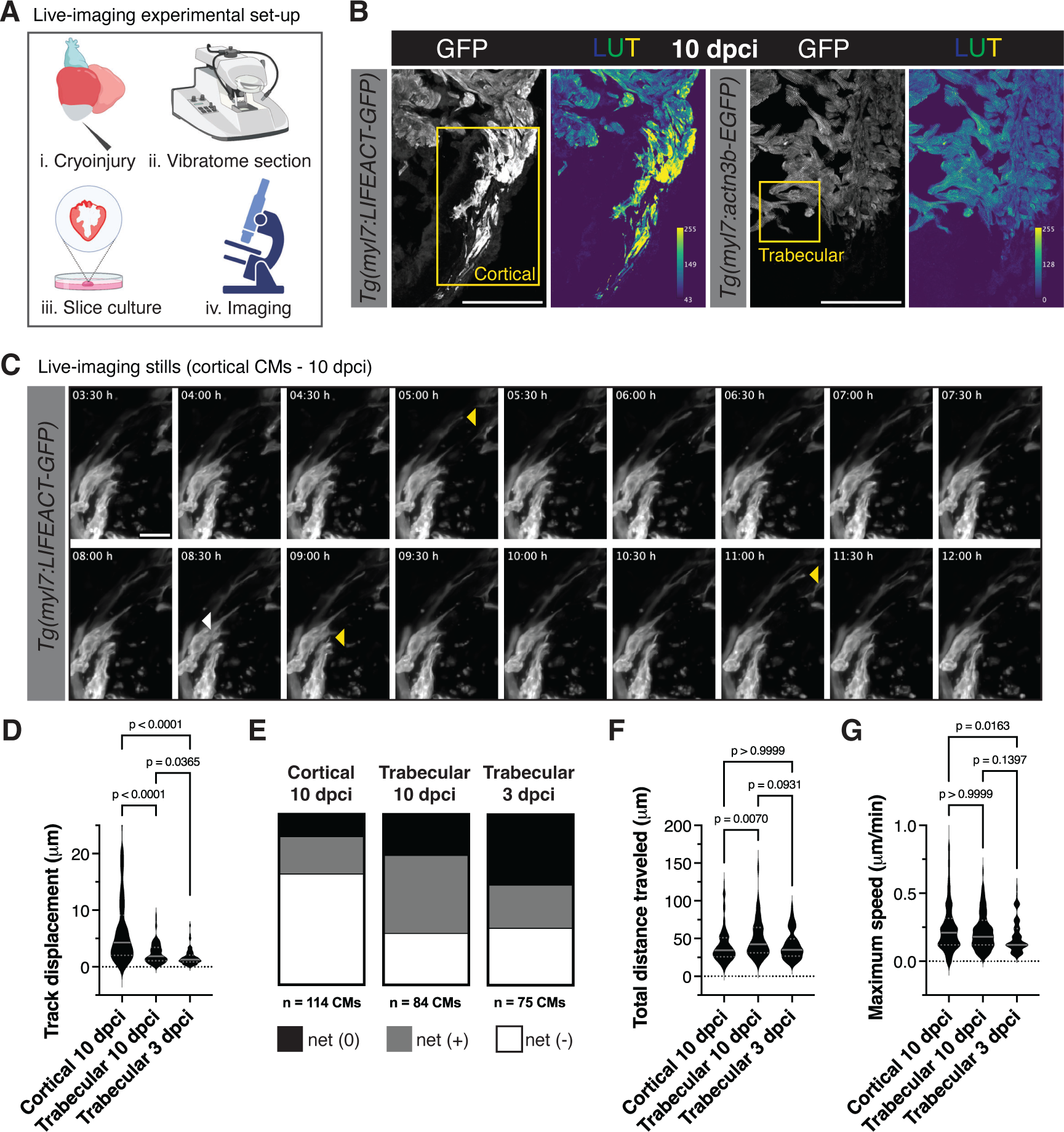
Live-imaging of CM invasion into the injured area during zebrafish heart regeneration. **(A)** Schematic of the experiment illustrating vibratome sectioning of cryoinjured hearts, slice culture, and live imaging. **(B)** GFP staining in thin sections of *Tg(myl7:LIFEACT-GFP)* (left) and *Tg(myl7:actn3b-EGFP)* (right) ventricles at 10 dpci. LUT (look-up table) images depict GFP intensity color coded according to the scale within the image. Yellow boxes denote the cortical/trabecular CM focus area of time-lapse imaging. **(C)** Time-lapse imaging of *Tg(myl7:LIFEACT-GFP)*+ cortical CMs at 10 dpci. Yellow arrowheads point to CM protrusions that display a net positive migration into the injured area and white arrowheads point to CM protrusions that display a net negative migration away from the injured area. **(D)** Quantification of track displacement from CM protrusions at the border zone in *Tg(myl7:LIFEACT-GFP)*+ cortical CMs at 10 dpci and *Tg(myl7:actn3b-EGFP)*+ trabecular CMs at 3 and 10 dpci. For cortical CMs, n=114 CMs were tracked from 6 ventricles; for trabecular CMs at 10 dpci, n=84 CMs were tracked from 4 ventricles; for trabecular CMs at 3 dpci, n=75 CMs were tracked from 5 ventricles. P-values were calculated using a Kruskal-Wallis test with Dunnett’s multiple comparisons test. **(E)** Distribution of tracked CM protrusions with a net positive displacement (> 1 μm into the injured area), net negative displacement (> 1 μm away from the injured area), or net zero displacement (< 1 μm) comparing t=0 and t=12 hours of live-imaging in *Tg(myl7:LIFEACT-GFP)*+ cortical CMs at 10 dpci and *Tg(myl7:actn3b-EGFP)*+ trabecular CMs at 3 and 10 dpci. **(F)** Quantification of total distance traveled by tracked CM protrusions in *Tg(myl7:LIFEACT-GFP)*+ cortical CMs at 10 dpci and *Tg(myl7:actn3b-EGFP)*+ trabecular CMs at 3 and 10 dpci. For cortical CMs, n=114 CMs were tracked from 6 ventricles; for trabecular CMs at 10 dpci, n=84 CMs were tracked from 4 ventricles; for trabecular CMs at 3 dpci, n=75 CMs were tracked from 5 ventricles. P-values were calculated using a Kruskal-Wallis test with Dunnett’s multiple comparisons test. **(G)** Maximum speed measured in tracked CM protrusions *Tg(myl7:LIFEACT-GFP)*+ cortical CMs at 10 dpci and *Tg(myl7:actn3b-EGFP)*+ trabecular CMs at 3 and 10 dpci. For cortical CMs, n=114 CMs were tracked from 6 ventricles; for trabecular CMs at 10 dpci, n=84 CMs were tracked from 4 ventricles; for trabecular CMs at 3 dpci, n=75 CMs were tracked from 5 ventricles. P-values were calculated using a Kruskal-Wallis test with Dunnett’s multiple comparisons test. Scale bars: 100 μm in **(B)**, 20 μm in **(C)**.

### Macrophages are associated with cardiomyocyte protrusions in regenerating zebrafish hearts

In the course of the live-imaging analysis above, we observed phagocytosed CM material in cells next to protruding cardiomyocytes. Using a *Tg(myl7:lck-mScarlet); Tg(mpeg1:EGFP)* double transgenic line, marking CM membrane and macrophages, respectively, we found that macrophages are in very close proximity to protruding CMs at the wound border zone (**Fig. 3A**). Quantification of macrophages 50 μm proximal and distal to the wound border revealed a peak in macrophage number at 7 and 10 dpci, corresponding to the peak in CM protrusion (**Fig. 3B**). To further understand the interaction of CMs and macrophages at the border zone, we performed time-lapse imaging of living ventricular sections from *Tg(mpeg1:EGFP); Tg(myl7:lck-mScarlet)* hearts at 10 dpci. Macrophages at the border zone extended multiple filopodia and were in close contact to CMs at the border zone (**Fig. 3C**, Supp. Fig. 3A, and **Supp. Video 5**). There was a striking difference in macrophage morphology in macrophages contacting CMs at the border zone and macrophages within the injured area (**Fig. 3C**). Macrophages within the injured area were significantly rounder (**Fig. 3D** and **Supp. Video 6**) and less protrusive. Previous studies have shown a correlation between macrophage morphology and phenotypic state, with anti-inflammatory or pro-healing macrophages displaying a more elongated morphology^40^. Together, these data suggest that macrophages in contact with border zone CMs are anti-inflammatory, or pro-healing, while those within the injured area are pro-inflammatory.

**Figure 3:**
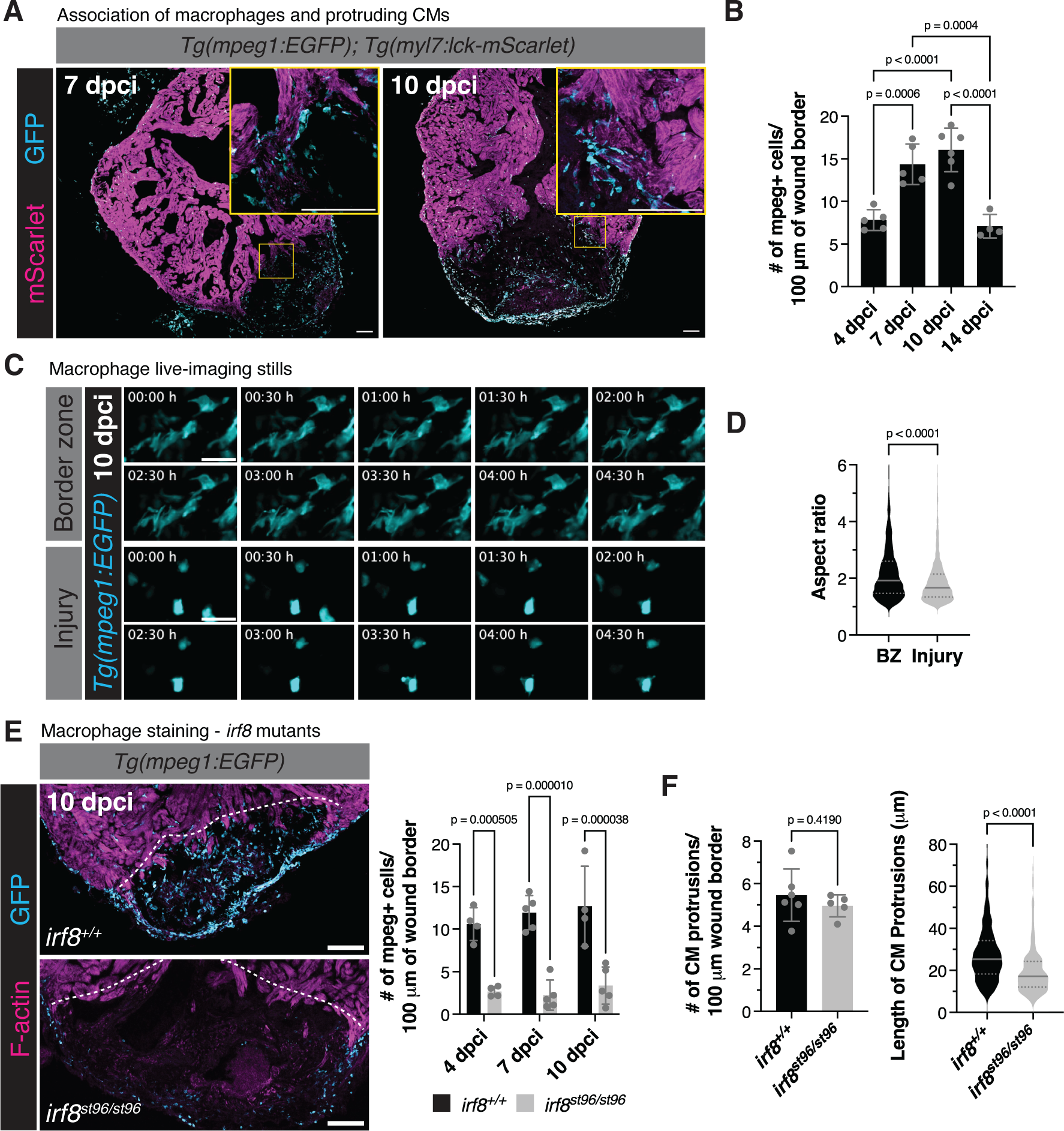
Macrophages are closely associated with invading CMs at the border zone. **(A)** Immunostaining of GFP and mScarlet in *Tg(mpeg1:EGFP); Tg(myl7:lck-mScarlet)* ventricles at 7 and 10 dpci. Yellow boxes denote the zoomed image from the wound border zone. **(B)** Quantification of the number of mpeg1:EGFP+ cells 50 μm proximal and distal to the wound border at 4 (n=5 ventricles), 7 (n=5 ventricles), 10 (n=6 ventricles), and 14 dpci (n=4 ventricles). P-values were calculated with multiple unpaired t-tests and corrected for multiple comparisons using the Bonferroni-Dunn method. **(C)** Time-lapse imaging of *Tg(mpeg1:EGFP)*+ cells at the border zone or within the injury at 10 dpci. **(D)** Quantification of aspect ratio of *Tg(mpeg1:EGFP)*+ cells at the border zone or within the injury at 10 dpci. n=1235 BZ macrophages and n=1267 macrophages within the injury were measured from 4 ventricles. P-values were calculated using a Mann-Whitney test. **(E)** Immunostaining of GFP and F-actin in *Tg(mpeg1:EGFP)* ventricles from *irf8^st96/st96^* mutants and wild-type siblings at 10 dpci (left). Quantification of *mpeg1*:EGFP+ cells 50 μm proximal and distal to the wound border at 4 (n= 4 ventricles), 7 (n=5 ventricles), and 10 dpci (n=4 wild-type and 5 *irf8* mutant ventricles) (right). P-values were calculated using multiple t-tests and were corrected for multiple comparisons using the Bonferroni-Dunn method. **(F)** Quantification of the number of CM protrusions per 100 micron of wound border (left) and length of CM protrusions (right) from thick cryosections of *irf8^st96/st96^* mutant (n=240 CMs from 5 ventricles) and wild-type sibling (n=485 CMs from 6 ventricles) ventricles at 10 dpci. P-values were calculated using an unpaired t-test (# of CM protrusions) and a Mann-Whitney test (CM protrusion length). Scale bars: 100 μm in **(A)** and **(E)**, 20 μm in **(C)**.

In order to determine whether macrophages contribute to the CM invasion process, we utilized *irf8^st96/st96^* mutant zebrafish^41^, which largely lack *mpeg1*+ macrophages in the heart at multiple timepoints following cryoinjury (**Fig. 3E**). Following cryoinjury in *irf8^st96/st96^* mutant zebrafish, we observed that while the number of CM protrusions was not affected at 10 dpci, the length of CM protrusions was significantly decreased in the absence of macrophages when compared to *irf8^+/+^* siblings (**Fig. 3F**). Because a lack of macrophages in the *irf8^st96/st96^* mutant hearts may alter cardiac physiology/homeostasis and the inflammatory response to cardiac injury^28, 42, 43^, we confirmed our results in a *Tg(mpeg1.1:NTR-YFP)* transgenic line, harboring macrophage-specific expression of the bacterial nitroreductase (NTR) enzyme. Treatment of adult fish with nifurpirinol, a nitroaromatic compound that was shown to be superior to metronidazole for NTR-mediated cell ablation^44^, from 4-6 dpci resulted in a large variability in macrophage ablation efficiency. However, we observed a significant correlation in the number of macrophages present at the wound border and the length in CM protrusions and no correlation between macrophage presence and number of CM protrusions, similar to the *irf8^st96/st96^* mutant (**Supp. Fig. 3B and 3C**).

### Resident macrophages are essential for ECM remodeling and cardiomyocyte invasion at the wound border zone

Based on our histological and live-imaging analyses, we hypothesized that the extracellular matrix (ECM) microenvironment adjacent to the border zone is important for CM protrusion/invasion and, further, that this ECM microenvironment is disrupted in the absence of macrophages. To address this hypothesis, we used a collagen hybridizing peptide (CHP) to visualize remodeling or degrading collagen at the border zone of regenerating hearts^45^. In *irf8^+/+^* wild-type ventricles, we observed collagen degradation/remodeling at the wound border zone (associated with protruding CMs) and extending into the wound. Strikingly, in *irf8^st96/st96^* mutant ventricles, there was a complete loss of CHP staining of collagen degradation/remodeling (**Fig. 4A**), suggesting that macrophage presence is necessary for ECM remodeling at the border zone and within the wound. As macrophages have previously been shown to deposit collagen in the wound area following cryoinjury in zebrafish hearts^30^, we stained *irf8^+/+^*wild-type and *irf8^st96/st96^* mutant ventricles with Picrosirius red to determine collagen content in the regenerating heart. We observed that collagen was localized at the wound border of *irf8^st96/st96^* mutant ventricles and at similar levels compared to *irf8^+/+^* wild-type ventricles (**Fig. 4B and 4C**), although the distribution of collagen differed slightly in the absence of macrophages. This difference in localization was likely due to the remaining presence of a large fibrin clot within the wound of *irf8^st96/st96^* mutant ventricles (**Supp. Fig. 4A**), suggesting that macrophage presence is essential for remodeling of the injured tissue following cardiac injury. This remodeling appears essential for organismal survival following cryoinjury, as a large majority of *irf8^st96/st96^* mutant fish died at mid-late stages of cardiac regeneration (**Supp. Fig. 4B**). As macrophages have also been shown to play a role in the activation of cardiac fibroblasts^45, 46^, we used quantitative reverse-transcription PCR (RT-qPCR) to analyze fibroblast gene expression in *irf8^+/+^* wild-type compared to *irf8^st96/st96^* mutant ventricles. We observed increased expression of canonical fibroblast markers, including *vim, acta2, col1a1a,* and *c4* in *irf8^st96/st96^* mutant ventricles compared to wild-type siblings at 10 dpci (**Supp. Fig. 4C**). Furthermore, we observed an increase in fibroblast genes that have been shown in previous studies to promote regeneration, including *fn1a, fn1b,* and *col12a1a* (**Supp. Fig. 4C**), suggesting that the lack of macrophages in *irf8^st96/st96^* mutants does not result in a decrease in fibroblast activation or expression of regenerative fibroblast marker genes. Altogether, these results indicate that macrophages are essential for collagen degradation/remodeling at the border zone, and that the lack of collagen degradation/remodeling in *irf8^st96/st96^* mutant ventricles is not due to the absence of collagen in the wound or a decrease in fibroblast gene expression programs.

**Figure 4:**
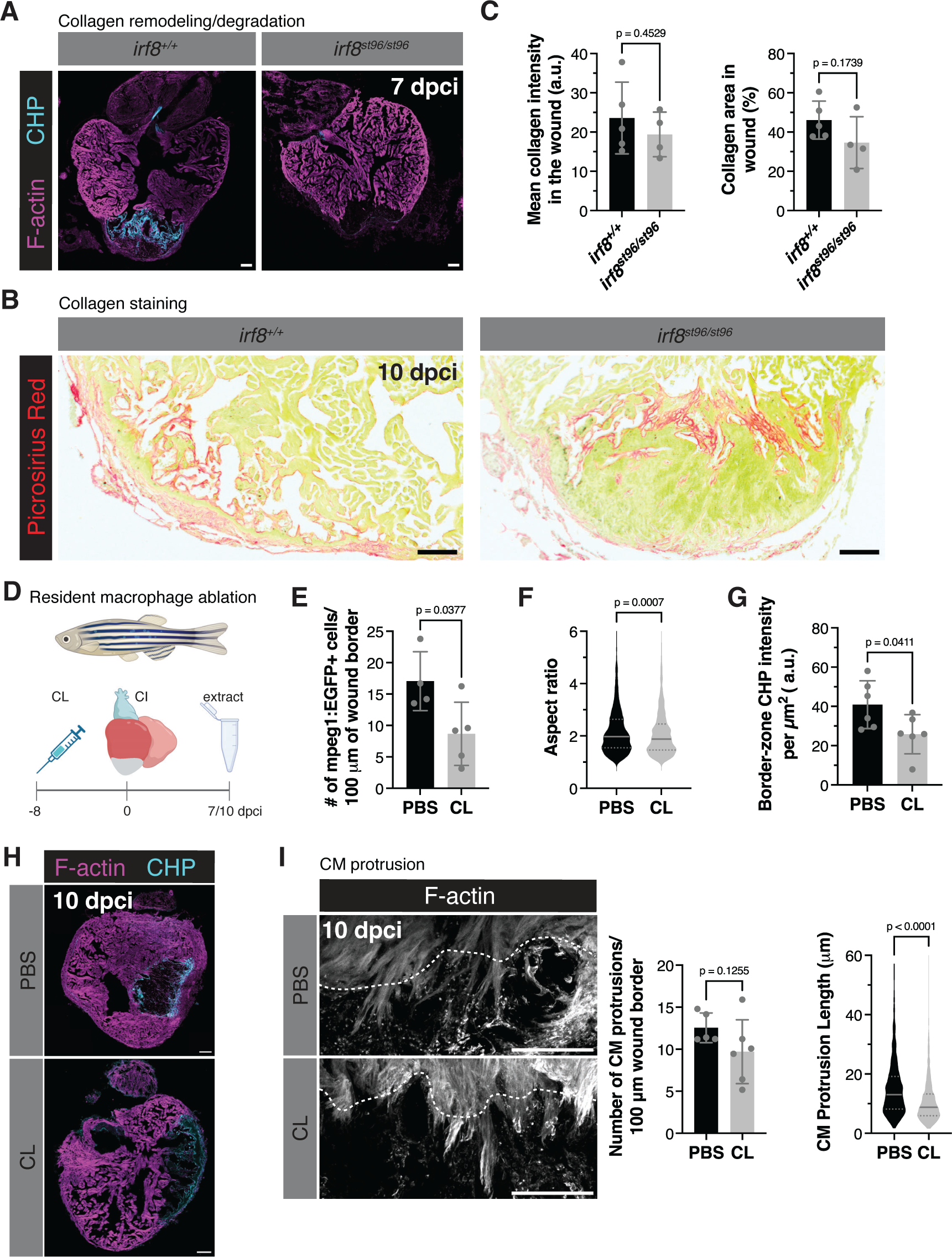
Macrophages are required for ECM remodeling at the wound border zone. **(A)** Collagen hybridizing peptide (CHP) and F-actin staining of *irf8^st96/st96^* mutant and wild-type sibling ventricles at 7 dpci. Images are representative of n=6 ventricles from each genotype. **(B)** Picrosirius red staining of collagen in *irf8^st96/st96^* mutant and wild-type sibling ventricles at 10 dpci. **(C)** Quantification of mean collagen intensity (arbitrary units, a.u.) and percentage of collagen area in the wound in *irf8^st96/st96^* mutant (n=4) and wild-type sibling (n=5) ventricles at 10 dpci. P-values were calculated using an unpaired t-test. **(D)** Schematic illustrating the experimental scheme to deplete resident macrophages with clodronate (CL) or PBS control liposomes 8 days prior to cryoinjury of ventricles. **(E)** Quantification of the number of *mpeg1*:EGFP+ cells 50 μm proximal and distal to the wound border in ventricles at 7 dpci in CL- (n=5) and PBS-liposome (n=4) injected fish. P-value was calculated using an unpaired t-test. **(F)** Quantification of aspect ratio of *Tg(mpeg1:EGFP)*+ cells 50 μm proximal and distal to the wound border in ventricles at 7 dpci in CL- (n=1087 BZ macrophages from 5 ventricles) and PBS-liposome (n=1497 BZ macrophages from 4 ventricles) injected fish. P-value was calculated using a Mann-Whitney test. **(G)** Quantification of CHP intensity at the border zone of 7 dpci ventricles in CL- (n=6) and PBS-liposome injected fish (n=6). P-values were calculated using an unpaired t-test. **(H)** Collagen hybridizing peptide (CHP) and F-actin staining of PBS control and CL-injected ventricles at 7 dpci. **(I)** Phalloidin staining of F-actin in thick cryosections of PBS liposome-injected and CL-injected zebrafish ventricles at 10 dpci (left). White dashed lines indicate the injury border. Quantification of the number of CM protrusions per 100 micron of wound border (middle) and length of CM protrusions (right) from thick cryosections of PBS liposome-injected (n=598 CMs from 5 ventricles) and CL-injected (n=845 CMs from 6 ventricles) zebrafish ventricles at 10 dpci. P-values were calculated using an unpaired t-test (# of CM protrusions) and a Mann-Whitney test (CM protrusion length).

Recently published studies have shown that resident macrophages are essential for the cardiac regenerative response^47^. In order to determine whether resident macrophages are important for ECM remodeling and CM protrusion, we administered clodronate liposomes (CL) 8 days prior to cryoinjury to deplete resident cardiac macrophages, performed cryoinjury, and determined the effect on ECM remodeling and CM protrusion at 7 and 10 dpci (**Fig. 4D**). Quantification of *Tg(mpeg1:EGFP)*+ cells at the wound border zone revealed that depletion of the resident macrophage population with CL resulted in a decrease of macrophages at the border zone (**Fig. 4E**). Furthermore, macrophages that were located at the border zone in resident macrophage-depleted hearts at 7 dpci exhibited a decrease in aspect ratio/elongation (**Fig. 4F**), suggesting that resident macrophages may contribute to the anti-inflammatory/pro-healing population of macrophages following cryoinjury. Notably, staining of remodeling/degrading collagen with CHP revealed a significant decrease in CHP intensity at the border zone in resident macrophage-depleted hearts at 7 dpci (**Fig. 4G** and **4H**). Accordingly, we observed no significant decrease in the number of CM protrusions, but a decrease in CM protrusion length at 10 dpci at the border zone (**Fig. 4I**), similar to *irf8^st96/st96^* mutant ventricles that largely lack macrophages. Altogether, these data indicate that macrophage presence, and in particular resident macrophage presence, at the wound border zone is essential for remodeling the ECM and CM protrusion/invasion.

### Single-cell RNA-sequencing analysis reveals border zone-specific gene expression signatures

In order to find potential regulators of CM invasion, we performed dissection of the wound border zone at 7 dpci and subjected these cells to single-cell RNA-sequencing (scRNA-seq, **Fig. 5A**). After removal of single cell transcriptomes that did not pass our quality control metrics (**Supp. Fig. 5A-5C**), 4596 cells remained. Dimensionality reduction with UMAP revealed 12 clusters of cells (**Fig. 5A**). Dot plots of unique markers for each cluster and for known cell type-specific marker genes verified the presence of known cell types at the border zone (**Supp. Fig. 5D-5E**). 36.3% of cells from our scRNA-seq dataset were CMs, including remote CMs (rCM) and border zone CMs (BZ CM), expressing previously described border zone marker genes, such as *mustn1b*, *tagln*, and *myl6*^48, 49^. 32.6% of cells from our scRNA-seq were fibroblasts, expressing known fibroblast genes, such as *col1a2, col1a1a, postnb,* and *fn1b*. We also observed specific fibroblast clusters, including Fb1 (*acta2*+), Fb2 (*c4* and *cxcl12a*+), a recently described regenerative Fb3 cluster (*col12a1a*+)^24^, and a small Fb4 (*fosab*+) cluster. 14.9% of cells from our scRNA-seq were endocardial/endothelial cells, including cluster EC1 (fibroblast-like EC (fEC) expressing endocardial markers, such as *fli1a* and *sele*, and fibroblast markers, such as *nppc* and *spock3*)^24^ and cluster EC2, expressing more traditional endothelial markers (eEC), including *kdrl* and *cldn5b*. The remaining 15.6% of cells from our scRNA-seq dataset were comprised of 4 immune/macrophage clusters, including mac1 (expressing *apoeb* and *wt1b*)^33^, mac2 (a small population expressing *defbl1*), mac3 (expressing canonical macrophage markers such as *mpeg1.1*, *mfap4*, *csfr1a, c1qa* and *grap2b*), and mac4 (ECM, expressing a number of macrophage markers, including *mpeg1.1* and *mfap4*, in addition to fibroblast-like genes, such as *postnb, col1a2, fn1b,* and *col12a1a*) (**Supp. Fig. 5D-5E**).

**Figure 5:**
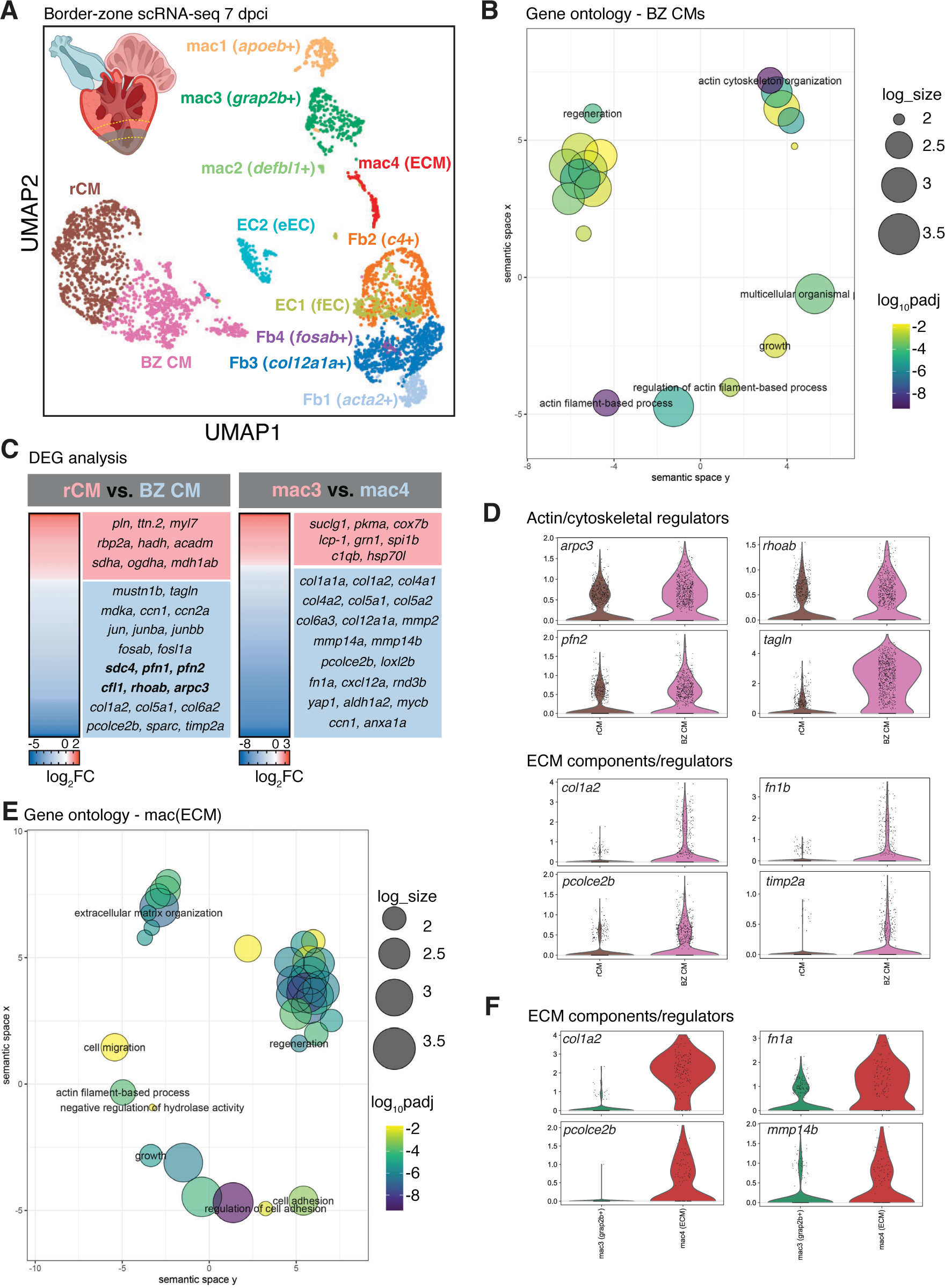
Single-cell RNA-sequencing reveals border zone-specific gene expression signatures in the regenerating heart. **(A)** UMAP plot of scRNA-seq from 4596 cells isolated from dissected border regions of regenerating zebrafish hearts at 7 dpci. Clusters were annotated by manual curation of known cells types at the border zone. The microdissected region of 7 dpci ventricles used for scRNA-seq is depicted in the upper left. **(B)** Gene ontology (GO) analysis of biological processes (BP) in differentially expressed genes enriched in BZ CMs compared to rCMs. log_10_padj was calculated in comparison to all genes in the zebrafish genome and log_size corresponds to the log_10_(number of annotations for the GO Term ID in zebrafish from the EBI GOA Database). **(C)** Differentially expressed genes (p<0.05) arising from pseudo-bulk comparisons of gene expression in CM versus BZ CMs (left) and mac3 versus mac4 (ECM, right) clusters. **(D)** Violin plots of expression of genes involved in actin cytoskeleton and ECM regulation in rCMs vs. BZ CMs clusters from scRNA-seq at 7 dpci. **(E)** Gene ontology (GO) analysis of biological processes (BP) in differentially expressed genes enriched in mac4(ECM) compared to mac3. log_10_padj was calculated in comparison to all genes in the zebrafish genome and log_size corresponds to the log_10_(number of annotations for the GO Term ID in zebrafish from the EBI GOA Database). **(F)** Violin plots of expression of genes involved in ECM composition/regulation in mac4 (ECM) vs. mac3 clusters from scRNA-seq at 7 dpci.

In order to investigate molecular mechanisms underlying CM invasion, we performed differential gene expression and gene ontology analyses between the BZ CM and remote CM clusters. As expected, differentially expressed genes (DEGs) that were higher in the remote CMs were enriched for biological processes such as ‘regulation of heart contraction’ and ‘generation of precursor metabolites and energy’, and included genes such as *myl7, pln,* and *ttn.2*, and genes encoding proteins involved in fatty acid oxidation (*rbp2a, hadh,* and *acadm*), cellular respiration and mitochondrial electron transport (*coq10b* and *sdha*), and the tricarboxylic acid (TCA) cycle (*mdh1ab*) (**Supp. Fig. 6A** and **Fig. 5C**). DEGs that were higher in border zone CMs included known border zone markers (*mustn1b* and *tagln*), along with genes that were enriched for biological processes such as ‘regeneration’ (*mdka, ccn1,* and *ccn2a*) and ‘actin cytoskeleton organization’ (*sdc4, pfn1, cfl1, rhoab,* and *arpc3*) (**Fig. 5B-5D** and **Supp. Fig. 6C**). Furthermore, we observed an enrichment of AP-1 transcription factor (TF) family members *jun, junba, junbb, fosab,* and *fosl1a* in border zone CMs, in line with our previously published data that AP-1 TFs are enriched in BZ CMs in response to cryoinjury^38^. Notably, Reactome pathway analysis of DEGs enriched in BZ CMs revealed pathways such as ‘cell:ECM interactions’, ‘collagen biosynthesis and modifying enzymes’, and ‘collagen degradation’ (**Supp. Fig. 6B**). The expression of genes, such as *col1a1a, col1a2, col6a2, pcolce2b, fn1a*, *itga5,* and *timp2a* were enriched in a subcluster of BZ CMs compared to remote CMs (**Fig. 5D** and **Supp. Fig. 6C**). Altogether, these data suggest that CMs upregulate gene expression programs that regulate actin cytoskeleton organization, cell:ECM interaction, and the ECM/collagen environment at the border zone to promote the migratory behavior we observe in our live-imaging data.

Based on our observations that macrophages are essential for CM protrusion length and for collagen degradation/remodeling at the border zone (**Fig. 3F** and **4A**), we focused our attention on the mac4 (ECM) cluster in our scRNA-seq dataset. Differential gene expression and gene ontology analyses between mac4 (ECM) and mac3 (expressing canonical macrophage markers) revealed an enrichment in biological processes such as ‘extracellular matrix organization’ (*col1a2, mmp14b, loxl2b, fn1a, col4a1,* and *col5a1*), ‘cell migration’ (*cxcl12a, rab13, cxcl18b, rnd3b,* and *fn1a*), and ‘regeneration’ (*yap1, aldh1a2, mycb, ccn1,* and *anxa1a*), and genes were enriched in the mac4 (ECM) cluster that were involved in ‘collagen degradation’ (*mmp2, mmp14a,* and *mmp14b*), and ‘collagen biosynthesis and modifying enzymes’ (*col12a1a, pcolce2b, col4a2,* and *col6a3*) (**Fig. 5C, 5E, 5F** and **Supp. Fig. 6D** and 6E). Taken together, our scRNA-seq analyses reveal a potential regulation of the ECM and ECM remodeling by border zone CMs and macrophages, which contributes to CM invasion into the injured cardiac tissue during regeneration.

### Mmp14b is a candidate regulator of extracellular matrix remodeling in border-zone cardiomyocytes and macrophages

We were intrigued by the presence of subclusters of CMs and macrophages at the border zone expressing several collagen genes and genes that are involved in collagen modification and degradation. Collagen hybridizing peptide (CHP) staining, marking remodeling/degrading collagen, and incubation of fresh sections with caged DQcollagen, fluorescently marking collagenase activity, revealed association of collagen degradation within the injured area, in line with previously published data^50^, and with border zone CMs at 10 dpci (**Fig. 6A** and **Supp. Fig. 7A**). Mining of previously published tomo-seq data from regenerating zebrafish hearts^48^ revealed an enrichment in *mmp14b* expression at the border zone, which we confirmed by *in situ* hybridization (**Fig. 6B**). Mmp14b is a membrane-tethered matrix metalloproteinase with multiple known ECM substrates^51^, including type I, II, and III collagens^52^, gelatin, fibronectin, and fibrin^53^, as well as non-ECM substrates, including pro-MMP2^54^, pro-MMP13^55^, and ADAM9^56^. Furthermore, MMP14 has been associated with cell migration in multiple contexts and has been observed in invadopodia, protrusive cellular structures found in invasive cells that promote cell attachment to and degradation of ECM^57^. *mmp14b* expression was upregulated in regenerating zebrafish hearts at 7 and 10 dpci (**Supp. Fig. 7B**) and enriched in multiple cell types from our scRNA-seq data, including BZ CMs, macrophages (mac1 and mac4(ECM)), endocardial cells (EC2), and fibroblasts (Fb2, Fb3, and Fb4) (**Fig. 6C** and **Supp. Fig. 7C**). Accordingly, sorting of CMs and macrophages by FACS revealed expression of *mmp14b* at 7 dpci and hybridization chain reaction (HCR) RNA-FISH revealed colocalization of *mmp14b* and CM and macrophage markers at the border zone at 10 dpci (**Supp. Fig. 7D** and 7E). Furthermore, we observed potential CM-specific and injury-responsive enhancer regions in intron 1 of the *mmp14b* locus from our previously published ATAC-seq data^38^, which differed from a recently described endothelial-specific *mmp14b* enhancer region from regenerating zebrafish hearts (**Supp. Fig. 7F**)^58^.

**Figure 6:**
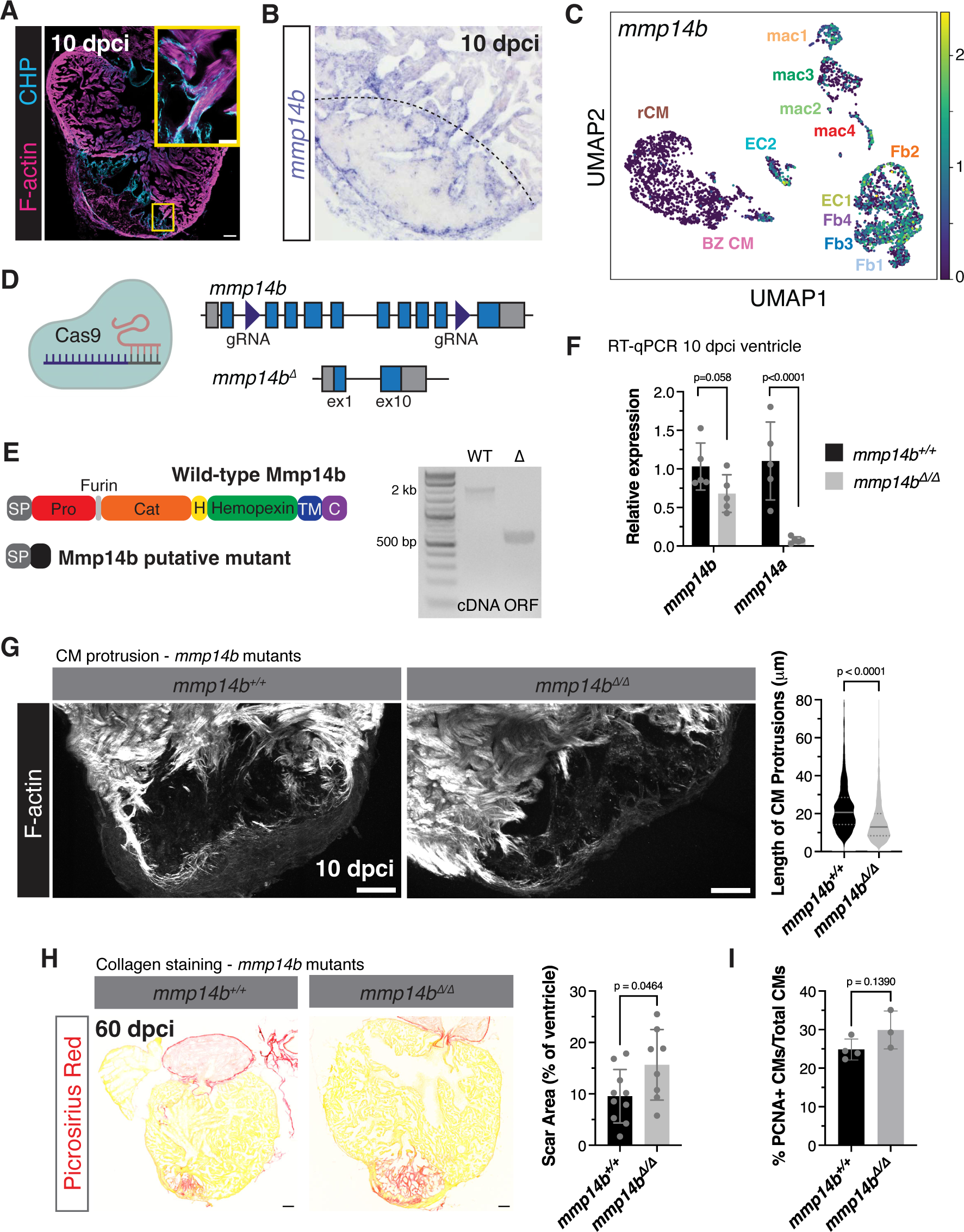
Mmp14b is a regulator of CM protrusion. **(A)** Collagen hybridizing peptide (CHP) and phalloidin staining of a wild-type ventricle at 10 dpci. Yellow box denotes the area in the zoomed image. **(B)** *in situ* hybridization of *mmp14b* expression in a wild-type ventricle at 10 dpci. Black dashed line denotes the approximate injury border. **(C)** UMAP plot of *mmp14b* expression from scRNA-seq of border zone cells at 7 dpci. **(D)** Schematic depicting the exon structure of the *mmp14b* locus and CRISPR/Cas9-induced full-length deletion between exons 2 and 9 of *mmp14b*. **(E)** Mmp14b wild-type and putative mutant protein domain structure (left). RT-PCR of the *mmp14b* open reading frame from wild-type and *mmp14b^11/11^* mutant embryos (right). SP, signal peptide; Pro, propeptide; Cat, catalytic domain; H, hinge region; TM, transmembrane domain; C, C-terminal tail. **(F)** RT-qPCR of *mmp14b* and *mmp14a* expression in single ventricles from *mmp14b^11/11^* (n=5) and wild-type siblings (n=5) at 10 dpci. P-values were calculated using an unpaired t-test. **(G)** Phalloidin staining of thick cryosections from *mmp14b^11/11^* and wild-type sibling ventricles at 10 dpci. Quantification of CM protrusion length (right) in n=1480 CMs from 9 ventricles (wild-type) and n=1124 CMs from 8 ventricles (*mmp14b^11/11^*). P-value was calculated using the Mann-Whitney test. **(H)** Picrosirius red staining of collagen in *mmp14b^11/11^* and wild-type sibling ventricles at 60 dpci (left). Quantification of scar area (% of ventricle area) in *mmp14b^11/11^* (n=8) and wild-type sibling (n=10) ventricles on the right. P-value was calculated using an unpaired t-test. **(I)** Quantification of CM proliferation within 100 μm of the wound border from PCNA/Mef2 immunostaining in *mmp14b^11/11^* (n=3) and wild-type sibling (n=4) ventricles at 7 dpci. P-value was calculated using an unpaired t-test. Scale bars: 100 μm, 20 μm in zoomed image in **(A)**.

We hypothesized that Mmp14b plays a role in remodeling collagen/the ECM environment at the border zone to regulate CM invasion. To test this hypothesis, we generated full locus deletion *mmp14b* mutants using CRISPR/Cas9 technology and 2 gRNAs flanking exon 2 and exon 9 of the *mmp14b* gene (**Fig. 6D**). Genotyping of larvae from a heterozygous *mmp14b^11/+^* incross led to Mendelian ratios of wild-type, heterozygous, and mutant larvae at 24 hpf (**Supp. Fig. 8A**). Deletion of *mmp14b* between exons 2-9 likely leads to an intact 5’ end of the Mmp14b protein with 29 amino acids (aa) containing the signal peptide and a frameshift consisting of 18 aa before a premature stop codon from exon 10 and presence of a shortened mutant *mmp14b* transcript in *mmp14b^11/11^* deletion mutant larvae was detected (**Fig. 6E**). We observed no gross morphological phenotypes during embryonic and larval development of *mmp14b^11/11^* mutants compared to wild-type siblings and *mmp14b^11/11^* mutants survived to adulthood. Based on previously published observations that single CRISPR/Cas9-induced mutations in *mmp14b* leads to nonsense mediated decay (NMD) and upregulation of the paralogous *mmp14a*^59^, we used RT-qPCR to assess levels of *mmp14a* and *mmp14b* in *mmp14b^11/11^* deletion mutants. In *mmp14b^11/11^* mutant hearts at 10 dpci, we observed a slight decrease in *mmp14b* mutant transcript levels, and *mmp14a* expression was significantly downregulated (**Fig. 6F**). This dramatic decrease in *mmp14a* levels in the *mmp14b^11/11^* mutants at 10 dpci was specific to the regenerating heart, as we only observed a slight decrease in *mmp14a* in *mmp14b^11/11^* mutants during larval stages (**Supp. Fig. 8B**). These results suggest that there is no genetic compensation from *mmp14a* in the *mmp14b^11/11^* full length deletion mutants.

### Mmp14b is an important regulator of ECM remodeling and cardiomyocyte invasion

In order to determine the importance of Mmp14b in CM invasion during cardiac regeneration, we used F-actin staining to compare CM protrusion number and length in *mmp14b^11/11^* full length deletion mutants at 10 dpci. We found that while there was no difference in the number of CM protrusions (**Supp. Fig. 8C**), we observed a significant decrease in the length of CM protrusions at the border zone in *mmp14b^11/11^* mutants when compared to wild-type siblings (**Fig. 6G**), similar to our observations in *irf8* mutant ventricles lacking macrophages. Using Picrosirius red staining, we found that *mmp14b^11/11^* mutants exhibited defects in cardiac regeneration, with the significant presence of collagen-containing scar tissue at 60 dpci when compared to wild-type siblings (**Fig. 6H**). These defects in cardiac regeneration were not due to differences in CM proliferation, as CM proliferation was not affected in the *mmp14b^11/11^* mutants generated here when compared to wild-type siblings at 7 dpci (**Fig. 6I**).

Due to the similarity in CM protrusion phenotypes between *mmp14b^11/11^* and *irf8^st96/st96^* mutants, we used collagen hybridizing peptide (CHP) staining to assess ECM degradation/remodeling at the border zone. Unlike *irf8^st96/st96^* mutants, *mmp14b^11/11^* mutants exhibited CHP staining within the wound at 10 dpci (**Fig. 7A**). However, quantification of CHP staining directly at the border zone revealed a significant decrease in CHP intensity in *mmp14b^11/11^* mutants compared to wild-type siblings at 10 dpci (**Fig. 7B**). Notably, this decrease in CHP intensity was accompanied by a corresponding decrease in *mpeg1:*EGFP+ cells at the border zone in *mmp14b^11/11^* mutants compared to wild-type siblings (**Fig. 7C and 7D**). These observations are in line with those from *irf8^st96/st96^* mutants, which lack macrophages, and with published data reporting a decrease in macrophage number proximal to the wound border at 3 days post apical resection in zebrafish hearts treated with a small molecule MMP14 inhibitor^60^. Taken together, these results indicate that Mmp14b is essential for macrophage presence and ECM degradation/remodeling at the wound border to promote CM invasion.

**Figure 7:**
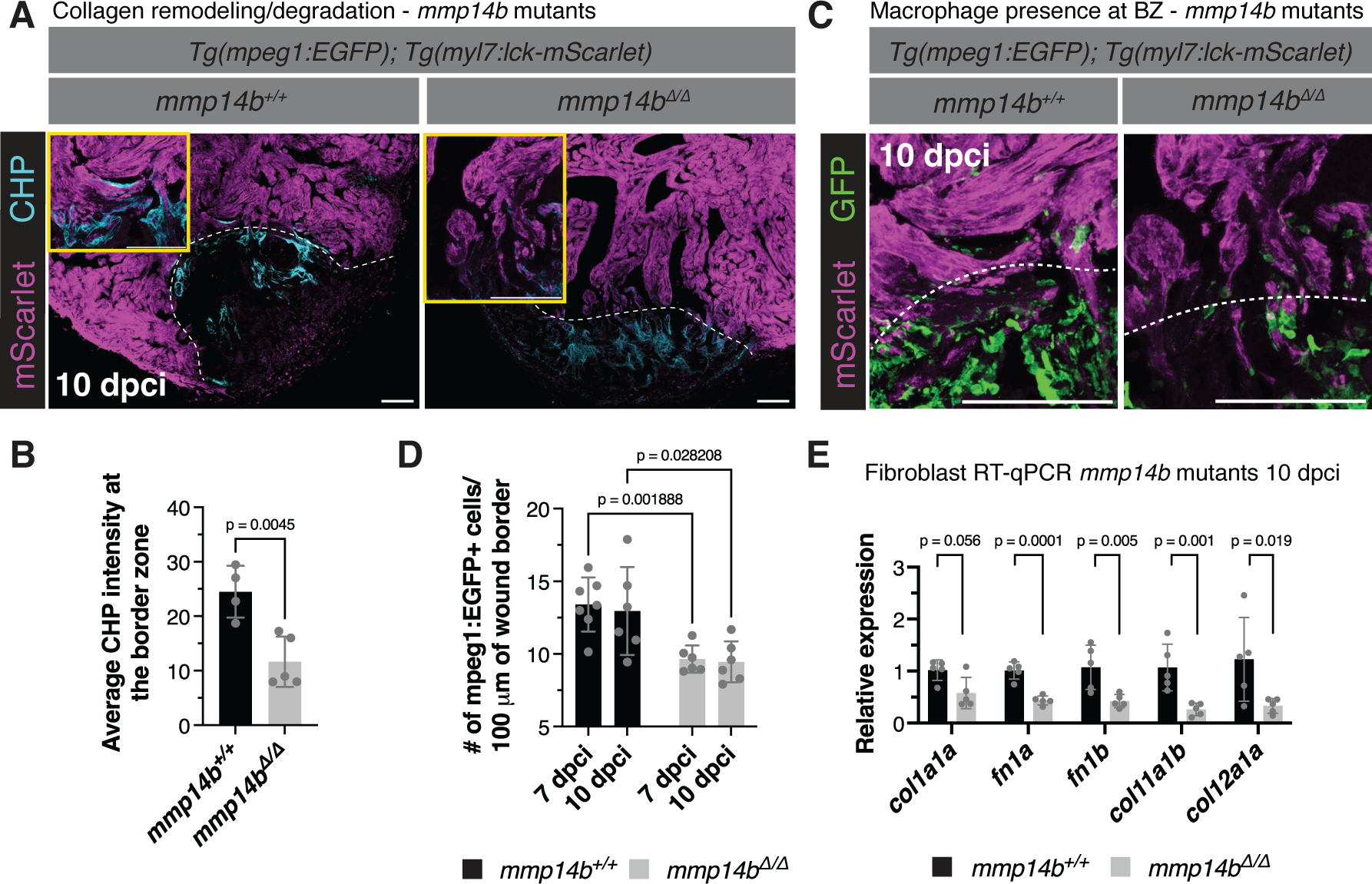
Mmp14b is essential for macrophage presence and ECM remodeling at the border zone. **(A)** Collagen hybridizing peptide (CHP) and mScarlet immunostaining in *Tg(mpeg1:EGFP); Tg(myl7:lck-mScarlet)* ventricles from *mmp14b^11/11^* and wild-type siblings at 10 dpci. White dashed lines indicate the wound border and yellow boxes contain zoomed images from the wound border zone. **(B)** Quantification of CHP intensity at the border zone (a.u., arbitrary units) in *mmp14b^11/11^* (n=5) and wild-type sibling (n=4) ventricles at 10 dpci.. P-value was calculated using an unpaired t-test. **(C)** GFP and mScarlet immunostaining in *Tg(mpeg1:EGFP); Tg(myl7:lck-mScarlet)* ventricles from *mmp14b^11/11^* and wild-type siblings at 10 dpci. White dashed lines indicate the approximate wound border. **(D)** Quantification of *mpeg1*:EGFP+ cells 50 μm proximal and distal to the wound border in ventricles from *mmp14b^11/11^* (n=6 ventricles at 7 and 10 dpci) and wild-type siblings (n=7 ventricles at 7 dpci and n=6 ventricles at 10 dpci). P-values were calculated using unpaired t-tests and corrected for multiple comparisons using the Holm-Sidak method. **(E)** RT-qPCR analysis of fibroblast marker genes in *mmp14b^11/11^* mutant (n=5) and wild-type sibling (n=5) ventricles at 10 dpci. P-values were calculated using an unpaired t-test or Mann-Whitney test (*col1a1a*). Scale bars: 100 μm.

As our scRNA-seq data revealed the expression of *mmp14b* in multiple cell types, including endothelial/endocardial cells and fibroblasts (**Supp. Fig. 7C**), we addressed the potential contribution of these cell types to the observed CM invasion defects in *mmp14b^11/11^* mutants. First, we performed immunostaining with an antibody for Aldh1a2, which has previously been shown to mark activated endocardium and epicardium^19^, in *mmp14b^11/11^* mutants and wild-type siblings at 7 dpci. We observed no significant difference in Aldh1a2 levels or expression pattern in both *mmp14b^11/11^* mutants, and *irf8^st96/st96^* mutants that largely lack macrophages, when compared to wild-type (**Supp. Fig. 9A**). We used RT-qPCR of a panel of fibroblast markers to determine whether *mmp14b* deletion affects fibroblast gene expression in response to cryoinjury. We found no significant differences in expression of canonical fibroblast markers, including *postnb, acta2,* and *vim*, or endocardial-derived fibroblast markers, *nppc* and *spock3,* in *mmp14b^11/11^* mutant compared to wild-type sibling ventricles at 10 dpci (**Supp. Fig. 9B**). However, we observed a significant decrease in expression of regenerative fibroblast markers, *col12a1a* and *col11a1b*, and ECM components, including *col1a1a, fn1a,* and *fn1b*, in *mmp14b^11/11^* mutant compared to wild-type sibling ventricles at 10 dpci (**Fig. 7E**). These data suggest that Mmp14b is important for the presence of a regenerative population of fibroblasts following cryoinjury. While these Col12a1a+ regenerative fibroblasts localize on the apical surface of the cryoinjured zebrafish heart at 7 dpci^24, 61^ and 10 dpci (**Supp. Fig. 9C**), and are therefore not in direct contact with trabecular protruding cardiomyocytes at the border zone, we cannot exclude fibroblast contribution to the *mmp14b* mutant phenotype that we observe.

### CM-specific *mmp14b* overexpression can promote CM protrusion

In order to determine whether CM-specific *mmp14b* overexpression is sufficient to promote CM invasion, we generated a transgenic line with spatial and temporal control of *mmp14b* overexpression (**Fig. 8A**). *Tg(hsp70l:loxP-EGFP-loxP-mmp14b-P2A-tagBFP); Tg(myl7:Cre-ERT2)* fish were treated with tamoxifen 2 days prior to cryoinjury (ethanol vehicle was used as a control) to allow for cardiomyocyte-specific *mmp14b* overexpression upon heat shock (**Fig. 8A and 8B**). Cardiomyocyte-specific *mmp14b* overexpression (hereafter referred to as CM:*mmp14b* OE) between 7 and 10 dpci led to an increase in the number of CM protrusions and length of those CM protrusions observed at the wound border zone when compared to control ventricles (**Fig. 8C**). Furthermore, Picrosirius red staining to visualize collagen and the wound area at 21 dpci revealed a trend towards decreased wound area in CM:*mmp14b* OE ventricles when *mmp14b* OE was induced from 7-20 dpci as compared to control (**Fig. 8D**). Notably, we observed that CM presence was increased on the apical surface of CM:*mmp14b* OE ventricles compared to control (**Fig. 8E**), in a similar location to where we observed highly protrusive cortically-located CMs in our live-imaging analysis (**Fig. 2**). Altogether, these observations indicate that CM-specific *mmp14b* overexpression is sufficient to induce CM protrusion and invasion of collagen-rich injured tissue after cardiac cryoinjury.

**Figure 8:**
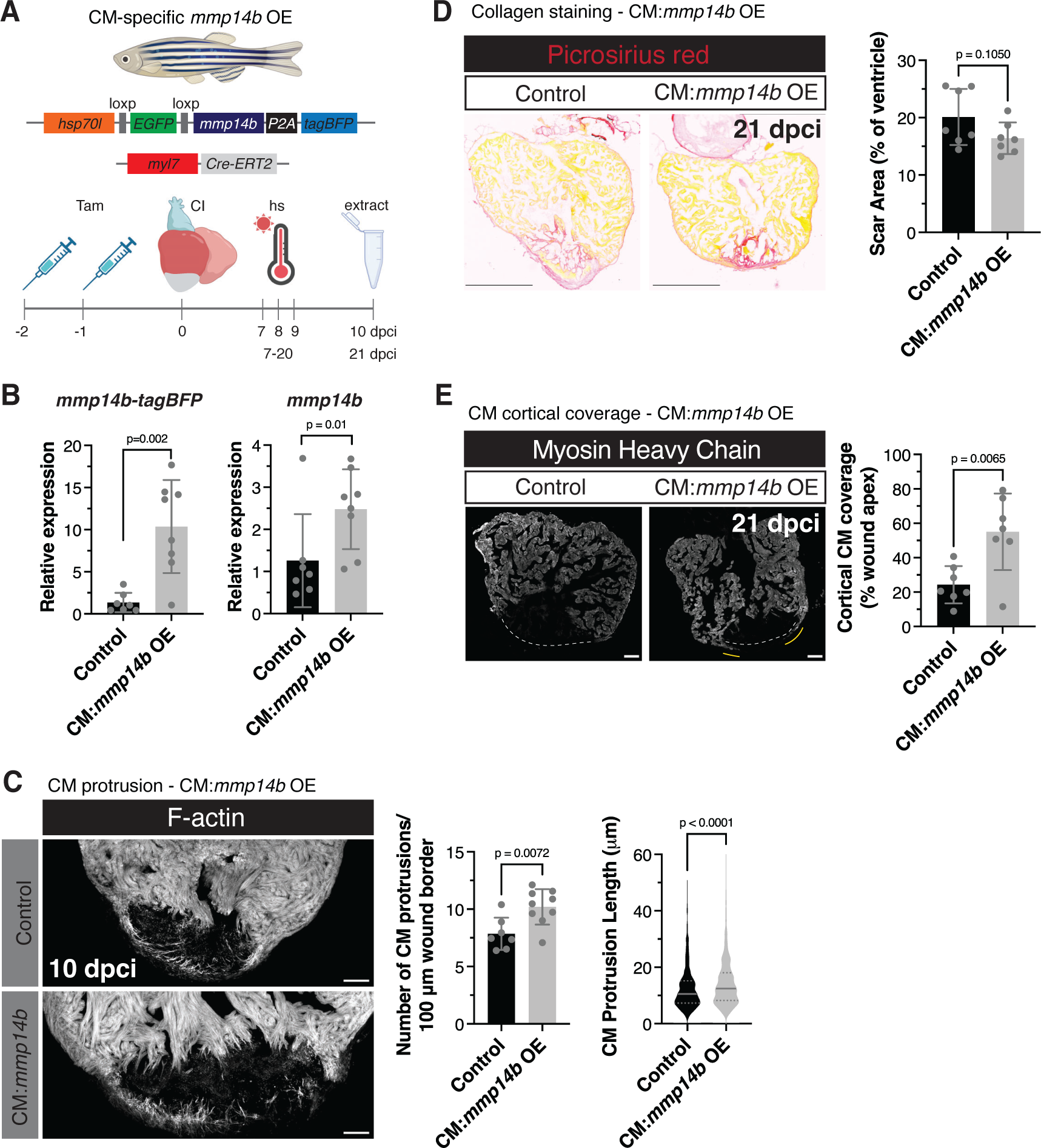
CM-specific overexpression of *mmp14b* promotes CM protrusion. **(A)** Schematic illustrating the *Tg(hsp70l:loxP-EGFP-loxP-mmp14b-P2A-tagBFPHA); Tg(myl7:Cre-ERT2)* line and experimental scheme to induce CM-specific overexpression of *mmp14b*. **(B)** RT-qPCR of *mmp14b-P2A-tagBFP* (left) and *mmp14b* (right) in control (n=7) and CM:*mmp14b* OE (n=8) zebrafish ventricles at 10 dpci. P-values were calculated using unpaired t-tests. **(C)** Phalloidin staining of F-actin in thick cryosections of control and CM:*mmp14b* OE zebrafish ventricles at 10 dpci. Quantification of the number of CM protrusions per 100 microns of wound border and length of CM protrusions from thick cryosections of control (n=814 CMs from 7 ventricles) and CM:*mmp14b* OE (n=1431 CMs from 9 ventricles) zebrafish ventricles at 10 dpci. P-values were calculated using an unpaired t-test (number of CM protrusions) or a Mann-Whitney test (CM protrusion length). **(D)** Picrosirius red staining of collagen in control and CM:*mmp14b* OE ventricles at 21 dpci (left). Quantification of scar area (% of ventricle area) on the right in control (n=7) and CM:*mmp14b* OE (n=7) ventricles. P-value was calculated using an unpaired t-test. **(E)** Myosin heavy chain immunostaining of control and CM:*mmp14b* OE ventricles at 21 dpci (left). White dashed lines denote the wound apex and yellow lines denote cortical CMs that have migrated over the wound apex. Quantification of cortical CM coverage (% wound apex) in control (n=7) and CM:*mmp14b* OE (n=7) ventricles at 21 dpci (right). P-values were calculated using an unpaired t-test. Scale bars: 100 μm in **(C)** and **(E)**, 500 μm in **(D)**.

## Discussion

Our data provide the first comprehensive analysis of CM protrusion into the injured collagenous tissue during the process of zebrafish heart regeneration. Through live-imaging of regenerating ventricular slices and scRNA-seq analyses, our results suggest that border zone CMs exhibit characteristics of actively migrating cells, displaying extension of motile filopodia directed towards the injured area and upregulation of gene expression programs that promote cell migration and remodel the ECM. We have previously published that AP-1 transcription factors are cell-autonomous regulators of CM protrusion and blocking AP-1 activity results in a dramatic reduction in CM protrusion at the wound border zone^38^. This cell-autonomous regulation is in line with studies from adult mouse models of cardiomyocyte regeneration that exhibit protrusion of CMs: in Hippo signaling-deficient mouse CMs, YAP binds to a number of genomic targets near genes that regulate actin cytoskeletal dynamics and cell:ECM interaction^10^, and CM-specific constitutively active *Erbb2* overexpression leads to the upregulation of both epithelial-to-mesenchymal transition (EMT) and ECM remodeling genes^15^. These data suggest that genetic programs that promote cell migration and invasion are upregulated in regenerating CMs and are important to replenish injured collagen-containing tissue with newly formed CMs.

We provide evidence that macrophages are closely associated with protruding/invading CMs and regulate the CM migration process through remodeling of the ECM at the border zone. Furthermore, through resident macrophage predepletion experiments, our data indicate that the resident macrophage population is at least partly responsible for this ECM remodeling process. Through scRNA-seq analysis of border zone cells, we find a population of macrophages that express genes important for remodeling and degradation of ECM, in line with recent published studies that independently corroborate the presence of this ECM macrophage cluster in the regenerating zebrafish heart^30, 62^. Notably, it has been recently shown that the resident macrophage population is essential for cardiac regeneration in zebrafish^29^, and that resident macrophage populations in the neonatal and adult mouse are essential for improving functional output following multiple types of cardiac injury and disease^63–65^. We suggest that one function of the resident macrophage population is to promote CM invasion and replenishment of collagen-containing injured tissue with CMs during regeneration. Likely, both macrophage-mediated remodeling of the ECM and CM-specific gene expression programs are necessary for efficient replenishment of the injured area with functional cardiomyocytes and it will be interesting to see if we can stimulate both processes in the nonregenerative mammalian heart to improve cardiac function following injury.

We show that Mmp14b is essential for macrophage presence and ECM remodeling at the wound border zone and that CM-specific *mmp14b* overexpression is sufficient to further promote CM invasion in regenerating zebrafish hearts. These observations are in line with a previously published study showing that MMP14 inhibition with a small molecule inhibitor leads to decreased macrophage presence proximal to the remaining cardiomyocytes in an apical resection model of cardiac injury in zebrafish^60^. A very recent study has shown that Mmp14b is an important regulator of scar resolution during zebrafish cardiac regeneration, in line with our data here^58^. However, the authors observed defects in CM proliferation that likely contribute to the defect in cardiac regeneration, which we do not observe with the *mmp14b* mutants generated here. This discrepancy may arise from differences in the mutation generated (deletion of exons 6-9 of *mmp14b*^58^ as opposed to deletion of exons 2-9 of *mmp14b* here) or differences in cell composition in the apical resection model compared to cryoinjury. While the authors show that *mmp14b* is expressed and plays an important role in endothelial cells following apical resection^58^, we find that *mmp14b* is expressed in a number of cell types following cryoinjury, including fibroblasts, endothelial/endocardial cells, macrophages, and cardiomyocytes. Our data also indicate that genetic deletion of *mmp14b* results in a decrease in the regenerative fibroblast markers *col12a1a* and *fn1a/fn1b*. These results raise multiple potential avenues of further study: for example, it is not yet known how regenerative fibroblasts contribute to CM protrusion and invasion of injured tissue. Moreover, previously published observations from immunostaining for Col12 in cryoinjured zebrafish hearts reveals localization of Col12 on the apical surface of the wound at 7 dpci and additional Col12 signal within the wound and neighboring border zone CMs at 14 dpci.^61^ Whether regenerative *col12a1a*+ fibroblasts migrate into the wound between 7 and 14 dpci and how Mmp14b contributes to this migration warrants further study.

Ultimately, we envision that our understanding of zebrafish CM invasion will aid the development of cell-based therapeutic strategies to promote engraftment of CMs into scar tissue to combat heart failure following myocardial infarction. Published studies have shown that while cardiomyocytes derived from human induced pluripotent stem cells (hiPSC) and human ventricular progenitor cells can improve cardiac functional output following cardiac injury, levels of CM engraftment into fibrotic scar tissue remain relatively low^66–68^. This phenomenon is likely due to multiple factors, including the lack of activation of gene expression programs necessary to stimulate CM invasion and differences in scar composition and stiffness between the regenerative zebrafish and nonregenerative mammalian heart. While fibroblasts are activated and deposit type I collagen in the cryoinjured zebrafish heart, this activation is transient and fibroblasts revert back to an inactivated state during the process of cardiac regeneration^22^. In the mammalian heart, while cardiac fibroblasts are essential for the initial wound healing response and for preventing ventricular rupture^69^, activation of fibroblasts, excessive ECM deposition, and the remaining presence of fibroblasts within the scar lead to a mature and permanent scar^70–72^. Published studies have revealed that decreasing the stiffness of the scar using small molecule compounds can extend the regenerative capacity of the neonatal mouse heart^73^, suggesting that the composition and stiffness of the scar limits regeneration or engraftment of cardiomyocytes. Therefore, it will likely be necessary to both promote invasion of exogenous cardiomyocytes and provide a permissive scar microenvironment when designing cell-based therapeutic strategies to repair the mammalian infarcted heart.

## Methods

### Zebrafish husbandry and lines

All zebrafish husbandry and experimentation were performed under standard conditions in accordance with institutional (MPG and Heidelberg University) and national (RP Darmstadt and RP Karlsruhe) ethical and animal welfare guidelines. Lines used in this study include: *Tg(myl7:LIFEACT-GFP)s974*^74^*, Tg(myl7:EGFP)twu26*^75^*, Tg(myl7:mVenus-gmnn)ncv43*^76^*, Tg(myl7:mCherry-cdt1)ncv68*^77^, *Tg(myl7:actn3b-EGFP)sd10*^78^*, Tg(gata4:EGFP)ae1*^79^*, Tg(mpeg1:EGFP)gl22*^80^*, Tg(mpeg1.1:NTR-YFP)w202*^81^*, irf8^st96^* ^41^, *Tg(cryaa:DsRed, −5.1myl7:CreERT2)pd10*^34^*, Tg(myl7:MKATE-CAAX)sd11Tg*^78^*, Tg(myl7:lck-mScarlet)bns561* (this study), *mmp14b^bns705^* (this study), and *Tg(hsp70l:loxP-EGFP-loxP-mmp14b-P2A-tagBFP)bns706* (this study). *irf8* mutant fish were genotyped using the following primers and high-resolution melt analysis (HRMA): 5’-ACGGCATACTAGTGAAGTAAAGGT-3’ and 5’-CTATAAGCCACTGTTTCAGTCTGC-3’.

### Generation of the *mmp14b* deletion line using CRISPR/Cas9

CRISPR/Cas9-mediated deletion of the *mmp14b* locus was performed using the Alt-R CRISPR-Cas9 system from Integrated DNA Technologies (IDT). CRISPR RNAs (crRNA) were designed using the Alt-R HDR Design Tool, were annealed to a transactivating crRNA (tracrRNA), and co-injected with Cas9 protein according to the “Zebrafish embryo microinjection” protocol from IDT (contributed by Jeffrey Essner, PhD). 2 gRNAs were co-injected to create a full locus deletion: 5’ gRNA targeting genomic sequence between exon 1 and 2 of *mmp14b* (5’-TACCCCCCAGTTCACCAACTTGG-3’) and exon 9 and 10 of *mmp14b* (5’-TATGACAGGACAATAATGAGAGG-3’). *mmp14b* deletion mutants were genotyped with the following primers: wild-type allele forward 5’-TGCATTCACACACATACACTGCGAC-3‘ and reverse 5’-ATTTACAACCACATCCCCCCTGCC-3’, and mutant allele forward 5’-AGACATAAGTGAAGAGTGAGAGAGG-3‘ and reverse 5’-TGGAGTCTTCGTTAGGGCAG-3‘.

### Generation of transgenic lines

The 1.5 kb fragment of the *hsp70l* promoter^82^ was PCR amplified and inserted into a pTol2 plasmid backbone. PCR amplification and conventional cloning methods were used to insert a floxed EGFP-stop cassette, the open reading frame of *mmp14b* (amplified from cDNA of 72 hpf zebrafish larvae), a P2A self-cleaving peptide, and HA-tagged tagBFP downstream of the *hsp70l* promoter. 75 pg of *Tol2* mRNA and 30 pg of construct were injected into one-cell stage AB embryos to generate the *Tg(hsp70l:loxP-EGFP-loxP-mmp14b-P2A-tagBFP)* line. For *Tg(myl7:lck-mScarlet)*, *lck* (ATGGGCTGCGTGTGCAGCAGCAACCCCGAG) was inserted upstream of the *mScarlet* ORF via PCR and *lck-mScarlet* was inserted into a pTol2 vector containing the −0.8 kb *myl7* promoter^83^. 75 pg of *Tol2* mRNA and 30 pg of construct were injected into one-cell stage AB embryos to generate the *Tg(myl7:lck-mScarlet)* line.

### Cryoinjury of zebrafish hearts and induction of transgene expression

Cryoinjury was performed according to previously published studies^5–7^. Briefly, adult male and female zebrafish between the ages of 4 and 12 months were anesthetized in 0.025% Tricaine and a small incision was cut in the skin and pericardial sac to expose the heart. The apex of the heart was touched with a liquid nitrogen-cooled metal probe and fish were transferred to a large beaker of system water for recovery.

For the induction of CM-specific *mmp14b* overexpression in *Tg(hsp70l:loxp-EGFP-loxP-mmp14b-P2A-tagBFP); Tg(myl7:CreERT2)* animals, adult zebrafish were anesthetized in 0.025% Tricaine and 0.5 mg/mL of 4-hydroxytamoxifen (4-HT) was injected intraperitoneally for 2 consecutive days. As a negative control, *Tg(hsp70l:loxp-EGFP-loxP-mmp14b-P2A-tagBFP); Tg(myl7:CreERT2)* adult zebrafish were injected with ethanol (EtOH) vehicle for 2 consecutive days. Following IP injection and cryoinjury, transgenic *Tg(hsp70l:loxp-EGFP-loxP-mmp14b-P2A-tagBFP); Tg(myl7:CreERT2)* zebrafish were incubated at 39°C for one hour daily to induce *mmp14b* overexpression.

### Immunostaining, confocal imaging, and quantification

Adult zebrafish hearts were extracted and fixed overnight in 4% (vol/vol) paraformaldehyde at 4°C, washed with 1x PBS, and cryopreserved in 30% (wt/vol) sucrose before embedding in O.C.T (Tissue-Tek) and stored at −80°C. 12 μm sections were collected for immunostaining analysis. For PCNA/Mef2 immunostaining, sections were washed twice with PBST (1x PBS, 0.1% Triton X-100), twice with dH_2_0, followed by permeabilization in 3% (vol/vol) H_2_O_2_ in methanol for 1 h at room temperature. Sections were washed twice in dH_2_0, twice in PBST, and then incubated in blocking solution (1x PBS, 2% (vol/vol) sheep serum, 0.2% Triton X-100, 1% DMSO) for 30 minutes to 1 h. Primary antibodies were incubated at 37°C for 3 hours, followed by three washes in PBST and incubation with secondary antibodies at room temperature for at least 1 h. Sections were washed in PBST and mounted in Vectashield Antifade mounting medium (Vector Labs). For collagen hybridization peptide (CHP), GFP, DsRed (mScarlet), mCherry, Myosin Heavy Chain (MHC), Col12a1a, and F-actin immunostaining, sections were washed twice with PBST, followed by permeabilization in PBSTx (1x PBS, 0.5% Triton X-100) for 0.5-1 h at room temperature, and incubated in blocking solution and then stained with primary antibodies in blocking solution (or CHP, heat dissociated at 80°C for 5 mins prior to dilution in blocking solution according to manufacturer’s instructions) overnight at 4°C. Washes and incubation with secondary antibodies were performed as above. For Aldh1a2 immunostaining, sections were washed twice with PBST and permeabilization was performed by incubation in acetone for 15 minutes at −20°C. Sections were then incubated in blocking solution and stained with primary antibodies in blocking solution overnight at 4°C. Washes and incubation with secondary antibodies were performed as above. Primary antibodies used in this study include: anti-PCNA (Abcam ab29) at 1:200, anti-Mef2 (Boster DZ01398-1) at 1:200, anti-GFP (Aves Labs GFP-1020) at 1:500, anti-DsRed (Takara 632496) at 1:300, anti-mCherry (Thermo Fisher M11217) at 1:200, anti-MYH1 (MHC, DSHB A4.1025) at 1:100, anti-Col12a1a (Boster DZ41260) at 1:200, anti-Aldh1a2 (Genetex GTX124302) at 1:100, collagen hybridizing peptide (CHP, 3Helix B-CHP) at 20 μM, and Alexa Fluor 647-Phalloidin (Thermo Fisher A22287) at 1:500. Alexa Fluor-coupled secondary antibodies were used (Thermo Fisher) at 1:500.

Imaging of immunostained sections was performed using a Zeiss LSM800 Observer confocal microscope with a 25x LD LCI Plan-Apochromat objective or a Leica Mica confocal microscope with a 20x HC PL FLUOTAR objective. Quantification of cardiomyocyte proliferation was performed in the area 100 μm proximal from the wound border in three nonconsecutive sections exhibiting the largest injured area from each heart (PCNA+Mef2+ CMs/total Mef2+ CMs). Quantification of CHP intensity was performed in the injured area 100 μm distal from the wound border in three nonconsecutive sections exhibiting the largest area from each heart. Quantification of Aldh1a2 intensity and % area was performed in the entire injury area in three nonconsecutive sections exhibiting the largest injured area from each heart. Quantification of cortical CM coverage in CM:*mmp14b* OE ventricles was performed by measuring the length of MHC+ signal along the apex of the wound and presented as the percentage of the length of the wound apex. Quantification of macrophage aspect ratio was performed in *mpeg1*:EGFP+ cells from thin, fixed cardiac tissue in the area 50 μm distal and 50 μm proximal to border zone CMs (BZ) or within the injured area (excluding the epicardium) in three nonconsecutive sections exhibiting the largest area from each heart. GFP signal was thresholded using the default algorithm and a binary image was used to quantify aspect ratio. All quantifications described above were performed in ImageJ.

To visualize cardiomyocyte protrusions at the border zone, wild-type hearts were fixed overnight in 4% PFA at 4°C and incubated in 30% sucrose/PBS overnight before embedding in Tissue-Tek OCT medium. 60 μm thick cryosections were obtained and allowed to dry at room temperature (RT) for 3 hours. Sections were then washed with 1x PBS twice at RT to remove OCT, permeabilized in 1x PBST (0.5% Triton X-100) for 2 hours, and incubated overnight with Alexa Fluor 647-Phalloidin (Thermo Fisher A22287) in blocking solution (1x PBS, 2% (vol/vol) sheep serum, 0.2% Triton X-100, 1% DMSO). Sections were washed 3x in PBST (0.1% Triton X-100) and embedded in Vectashield Antifade mounting media (Vector Labs). Thick sections were imaged using a Zeiss LSM800 Observer confocal microscope with a 25x LD LCI Plan-Apochromat objective or a Leica Mica confocal microscope with a 20x HC PL FLUOTAR objective. Cardiomyocyte protrusions were quantified using a maximum intensity projection image from at least 2 sections exhibiting the largest injured area and containing largely trabecular cardiomyocytes. Quantification of protrusions into the injured area and measurement of length were performed in ImageJ.

### Histological staining, imaging, and quantification

Adult zebrafish hearts collected for histology at 10 dpci were extracted and fixed overnight in 10% formalin at 4°C, washed with PBS, dehydrated in ethanol gradient, washed in Xylene followed by 50% Xylene/ 50% paraffin, and embedded in paraffin. 5 μm sections were obtained for histological analysis. Adult zebrafish hearts collected for histology at 21, 30, and 60 dpci were extracted and fixed overnight in 4% paraformaldehyde at 4°C, washed with PBS, cryopreserved in 30% sucrose and embedded in O.C.T. (Tissue-Tek) and stored at −20°C. 12 μm sections were obtained for histological analysis.

Picrosirius Red staining was performed using the Picrosirius Red stain (Morphisto). FFPE sections were incubated in Bouin’s solution overnight at room temperature, followed by a rinse in running water for 10 minutes. Incubation with Picrosirius Red solution and subsequent steps were performed according to manufacturer’s instructions. Cryosections were post-fixed 2 min in 4% paraformaldehyde at room temperature, followed by overnight incubation in Bouin’s solution at room temperature and washed in running water for 10 minutes. Sections were incubated in Picrosirius Red solution for 90 min at room temperature and subsequent steps were performed according to manufacturer’s instructions. Sections were mounted in Entellan (Sigma).

AFOG staining was performed using the Acid Fuchsin Orange G Kit (Biognost). FFPE sections were incubated in Bouin’s solution overnight at room temperature, followed by a rinse in running water for 10 minutes. Subsequent steps were performed according to manufacturer’s instructions. Sections were mounted in Entellan (Sigma).

Imaging of histological staining was performed with the Leica Mica using widefield imaging, with a 10x or 20x HC PL FLUOTAR objectives. Scar area was quantified from Picrosirius Red staining of three to five nonconsecutive sections from each heart relative to the ventricle area in ImageJ.

### *In situ* hybridization

*In situ* hybridization from ventricle sections was performed on cryosections according to standard procedures. Briefly, sections were dried at 50°C, fixed in 4% (vol/vol) paraformaldehyde, treated with 5 μg/mL Proteinase K (Roche) and post-fixed in 4% (vol/vol) paraformaldehyde. Acetylation of sections was performed by incubation in 0.1 M triethanolamine/0.25% acetic anhydride with stirring, followed by washing in PBS, and prehybridization at 70°C in hybridization buffer (50% formamide, 5x SSC, 0.1% Tween-20, 50 μg/mL heparin, 500 μg/mL yeast tRNA, pH 6.0). Digoxigenin-labeled RNA probes (200 ng/mL) were hybridized to tissue sections overnight at 70°C in a hybridization oven. Washes with 2x SSC and 0.2x SSC were performed at 70°C, followed by washes in PBT (1x PBS, 2 mg/mL BSA, 0.1% Triton X-100) and blocking in PBT/10% inactivated sheep serum. Sections were incubated overnight at 4°C with Anti-Digoxigenin-AP antibody (1:2000, Roche) in blocking solution. Sections were washed in PBT and alkaline phosphatase buffer and stained with BM Purple (Roche) staining solution. The following primers were used to amplify PCR products which were used to generate in situ hybridization probes with T7 RNA polymerase (Promega) and Digoxigenin-UTP (DIG RNA Labeling Mix, Roche): *mmp14b* Fwd: 5’-AATTAACCCTCACTAAAGGGAGAATCTGGAGGACACCCTCGAC-3’, Rev: 5’-TAATACGACTCACTATAGGGAGAGGCAGCCCATCCAATCCTTA-3’. Imaging of *in situ* hybridized sections was performed using a Nikon SMZ25 stereomicroscope.

### Hybridization chain reaction staining

mRNA expression of *mmp14b* was verified by HCR^TM^ RNA fluorescence *in situ* hybridization (RNA-FISH). Ventricular tissue was pretreated with methanol at −20°C overnight before embedding in OCT. GFP/DsRed immunostaining was performed according to the aforementioned protocol before proceeding with FISH following the manufacturer’s protocol (Molecular Instruments). Notably, during the hybridization with 20nM *mmp14b* probe, sections were incubated overnight in dark humid section box at 37°C, without being covered by cover slips.

### DQCollagen staining

DQCollagen staining was performed according to Gamba et al.^50^ with the following alterations. Briefly, unfixed cryoinjured hearts were incubated overnight at 4°C in 30% sucrose/PBS and embedded in OCT medium (Tissue-Tek). 12 μm cryosections were obtained and embedding medium was removed by washes in PBST (0.1% Triton X-100). Sections were preincubated for 5 minutes at RT in 1x reaction buffer (50 mM Tris-HCl, 150 mM NaCl, 5 mM CaCl_2_, 0.2 mM sodium azide, pH 7.6) and then incubated in 0.1 mg/mL DQCollagen (fluorescein conjugate, Thermo Fisher D12060) in 1x reaction buffer for 1 hour at RT. Sections were washed 3x in 1x PBST (0.1% Triton X-100) and incubated in Alexa Fluor 647-coupled phalloidin in 1x PBST (0.1% Triton X-100) for 1 hour at RT. Sections were washed 3x in 1x PBST (0.1% Triton X-100), mounted in Vectashield Antifade mounting medium (Vector Labs), and imaged with confocal microscopy as described above for immunostained sections.

### Live-imaging analysis of ventricular tissue slices

Zebrafish euthanasia and heart extraction were performed following previous reports^37^. After removal of blood and stopping the heartbeat, hearts were oriented with the former site of atrium facing up and mounted in 2.5% low EEO agarose (Sigma-Aldrich #A0576) dissolved in LAF medium [Leibovitz’s L15-Medium with GlutaMAX™ (Gibco #31415029)/1× MEM Non-Essential Amino Acids Solution (Gibco #11140035)/20 mM 2,3-Butanedione monoxime (Sigma-Aldrich #B0753)/10% Fetal Bovine Serum (Gibco #A3160401)/100 μg/ml Primocin (InvivoGen #ant-pm-1)/1% Pen/Strep (Gibco #15070063)] in a cryomold. After solidifying for 5 minutes, agarose blocks were sectioned using a vibratome [Precisionary Instruments #VF-310-0Z, slice thickness =100 μm, speed = 1.8, oscillation frequency = 5] and collected in LAF medium. Sections were transferred to μ-Slide 8-Well High ibiTreat plate (ibidi #80806) precoated with 11 μg/cm^2^ fibronectin (Sigma-Aldrich #F0895) containing LAF medium. The medium was largely removed to allow sections to adhere to the plate during incubation at 28.5°C for 90 minutes. LAF medium was then supplied back to the plate storing at 28.5°C before imaging.

The ibiTreat 8-well plate was directly used for time-lapse imaging using a Nikon AX confocal microscope. For Tg(*myl7*:*LIFEACT-EGFP; myl7:nls-DsRed*) and Tg(*myl7*:*actn3b-EGFP; myl7:nls-DsRed*), imaging was performed with a 1.5 μm *z*-step size and a 15-minute interval using a 20× objective for 12 hours. For *Tg(myl7*:*lck-mScarlet; mpeg1:EGFP*), imaging was performed with a 1.5 μm *z*-step size and a 15-minute interval using a 20× objective for 6 hours. Time-lapse videos were corrected for drift using the StackReg plug-in in ImageJ. Cardiomyocyte protrusion ends were tracked manually, and migratory parameters were measured using the TrackMate plug-in^84^ in ImageJ.

### Macrophage ablation

Ventricles from *Tg(mpeg:NTR-YFP)* adult fish were cryoinjured as described above. Cryoinjured fish were then incubated in a 5 μm nifurpirinol or DMSO-control water bath from 4-6 days following cryoinjury (replenished daily). Hearts were extracted at 7 dpci and subjected to immunostaining with GFP antibody to check macrophage ablation efficiency and phalloidin staining to quantify CM protrusion at the border zone as described above.

To ablate resident macrophages, *Tg(mpeg1:EGFP)* adult fish were IP injected with 10 μl clodronate liposomes (5 mg/ml) (Liposoma, Amsterdam, The Netherlands) or PBS 8 days prior to cryoinjury. Hearts were extracted at 7 or 10 dpci followed by cryosection, CHP staining, and phalloidin staining, as described above. Quantification of macrophage aspect ratio, CHP intensity and CM protrusion at the border zone were performed as described above.

### Single-cell RNA-sequencing of border zone cells

Zebrafish hearts were cryoinjured as above and extracted at 7 dpci. The border zone was roughly dissected and a pool of border zone cells from 8 hearts was dissociated using the Pierce Primary Cardiomyocyte Isolation kit (Thermo Fisher 88281) according to manufacturer instructions. The cell suspensions were depleted for dead cells using a LeviCell 1.0 device (Levitas Bio) and were counted with a Moxi cell counter and diluted according to manufacturer’s protocol to obtain 10,000 single cell data points per sample. Each sample was run separately on a lane in Chromium controller with Chromium Next GEM Single Cell 3ʹ Reagent Kits v3.1 (10xGenomics). Single-cell RNA-seq library preparation was done using standard protocols. Sequencing was done on a Nextseq2000 and raw reads were aligned against the zebrafish genome (DanRer11) and counted by StarSolo^85^, followed by secondary analysis in Annotated Data Format. Preprocessed counts were further analyzed using Scanpy^86^. Basic cell quality control was conducted by taking the number of detected genes and mitochondrial content into consideration. We removed 23 cells in total that did not express more than 300 genes or had a mitochondrial content greater than 40%. Furthermore, we filtered 9874 genes if they were detected in less than 30 cells (<0.01%). Raw counts per cell were normalized to the median count over all cells and transformed into log space to stabilize variance. We initially reduced dimensionality of the dataset using PCA, retaining 50 principal components. Subsequent steps, like low-dimensional UMAP embedding (McInnes & Healy, https://arxiv.org/abs/1802.03426) and cell clustering via community detection (Traag et al., https://arxiv.org/abs/1810.08473), were based on the initial PCA. Final data visualization was done by Scanpy and CellxGene packages (DOI 10.5281/zenodo.3235020). Differential gene expression analysis (DEG) was performed using Scanpy. Gene ontology analysis of DEGs was performed using Panther (v18.0)^87^ and visualized using REVIGO^88^.

### Gene expression analysis by RT-qPCR

RNA from adult zebrafish ventricles and larvae was isolated by resuspension of cells in TRIzol (Thermo Fisher), addition of chloroform, and centrifugation according to manufacturer’s instructions. RNA from the aqueous layer was purified using the RNA Clean and Concentrator Kit (Zymo). Total RNA was reverse transcribed to cDNA using the iScript Advanced cDNA kit (Biorad) according to manufacturer’s instructions. qPCR was performed using SsoAdvanced Universal SYBR Green Supermix (Biorad) and the LightCycler 480II (Roche). The following primers were used for gene expression analysis by RT-qPCR: *rpl13* 5’-TCTGGAGGACTGTAAGAGGTATGC-3’ and 5’-AGACGCACAATCTTGAGAGCAG-3’, *mmp14b* 5’-GGAAAATGATCTGGAGCGGGTT-3’ and 5’-AGCCATGCCTCAGGTTTCATA-3’, *mmp14b_3’UTR* 5’-GCGCAGTTACCAAATGCACA-3’ and 5’-CATGGGGTGAGAATGGACCC-3’, *mmp14a* 5’-cttcagagtttgggaggccg-3‘ and 5’- gtacgcatgggccaggaaac-3’, *fn1b* 5’-GCCATTGGTACTGGATCTGCAG-3’ and 5’- GTTCTGATCCAGCTCATGCCATTG-3’, *fn1a* 5’-AAACTTGGAGCGGCTGCGG-3’ and 5’- CACAGGTGCGATTGAACACG-3’, *postnb* 5’-GACCCGAGTTATCCAGGGAGAG-3’ and 5’- CTCAATCACACGAGTGACCTTGG-3’, *fap* 5’-TTCGCTTGGAGTGGGCGAC-3’ and 5’- TTCATCCCCCTGGAAACCACTTC-3’, *vim* 5’-TCCAGCCGGCAGTACAGCAG-3’ and 5’- CTCAAGCCTTTACTCGCGTAC-3’, *col1a1a* 5’-GCCAGGTCTACAATGACAGGG-3’ and 5’- ATCACTTCGTCGCACATTACGG-3’, *col11a1b* 5’-CCCAAAAGGCCCACCTGGG-3‘ and 5’- TGGAAACCAGTCTCTCCTCGC-3’, *acta2* 5’-CTCTGGAGAAGAGCTATGAGCTTC-3’ and 5’-TCCCGATGAAGGACGGCTG-3’, *col12a1a* 5’-TTGACTCTAGTACTCAGTTCAGTAG-3’ and 5’-TTCCTCCATGACATTTGCACTGTG-3’, *nppc* 5’-GGAGCTGTCTGAGTTCCTGG-3’ and 5’-TGTTCGTTCATGATCCGCGC-3’, *spock3* 5’-AGAGATGAGGTGGAGGAACCAG-3’ and 5’-ACATTTTATCTTCAGACATGGATCTTTGG-3’, *c4* 5’- TTCATATGTTCAACTTGCTGCC-3’ and 5’-CCCCGTTTTGTTGACAAATAGATG-3’.

### Sorting of cells by FACS for gene expression analysis

*Tg(myl7:mKATE-CAAX); Tg(mpeg1:EGFP)* adult fish were cryoinjured and ventricles extracted at 7 dpci. Tissue was dissociated using the Pierce Primary Cardiomyocyte Isolation kit (Thermo Fisher 88281) according to manufacturer’s instructions and single cells were resuspended in DMEM/10% FBS/1x glutamate. mKATE+ CMs, EGFP+ macrophages, and mKATE-EGFP-cells were sorted using a FACS Aria III and subjected to gene expression analyses as above.

### Statistical analysis

For all bar graphs, each point represents measurements from a single ventricle, the bar depicts the mean, and error bars depict standard deviation. For all violin plots, the center line depicts the median and the dotted lines depict the 1^st^ and 3^rd^ quartiles. All statistical analyses for experimental data were performed in GraphPad Prism. Distribution of data from all sample groups was assessed using the Shapiro-Wilks normality test. Comparative statistics between two sample groups was performed using the unpaired t-test for parametric data or the Mann-Whitney test for nonparametric data. Comparative statistics between more than two sample groups was performed using ordinary one-way ANOVA for parametric data or the Kruskal-Wallis test for nonparametric data. Specific multiple comparison tests used for experimental data with more than two sample groups can be found in the figure legends.

## Data Availability

Single-cell RNA-seq data from this study have been deposited in the Gene Expression Omnibus (GEO) database under the accession number: GSE251856

## Acknowledgements

We thank Carmen Buettner and Sara Schumann for excellent technical assistance, and Ann Atzberger and Kikhi Khrievono for assistance with FACS. Time-lapse imaging was performed at the Nikon Imaging Center (Heidelberg University) and we are grateful to Ulrike Engel and Christian Ackermann for their assistance. We thank members of the Stainier, Beisaw, and van den Hoogenhof labs for discussion and input, and Felix Gunawan and Chi-Chung Wu for critical feedback on the manuscript. This publication was supported through state funds approved by the State of Baden-Wuerttemberg for the Innovation Campus Health and Life Science Alliance Heidelberg Mannheim (A.B.), the German Center for Cardiovascular Research ((DZHK, Project number 81X2200317), A.B. and D.Y.R.S.), and the Max Planck Society (D.Y.R.S). F.C. is supported by an Inter-institutional Postdoctoral Fellowship from the Health and Life Science Alliance Heidelberg Mannheim, and A.B. was supported by the NHLBI (NIH) under award number 1F32HL143839.

## Author Contributions

Conceptualization: A.B.; Methodology: R.P.; Formal Analysis: S.G.; Investigation: F.C., B.W., K-H.W., I-T.L., J.D., T.L., A.B.; Writing – Original Draft: A.B.; Writing – Review and Editing: F.C., B.W., S-L.L, D.Y.R.S.; Visualization: F.C., B.W., A.B.; Supervision: S-L.L, D.Y.R.S., A.B.; Funding acquisition: D.Y.R.S., A.B.

## Supplementary Figures

**Supp. Figure 1:**
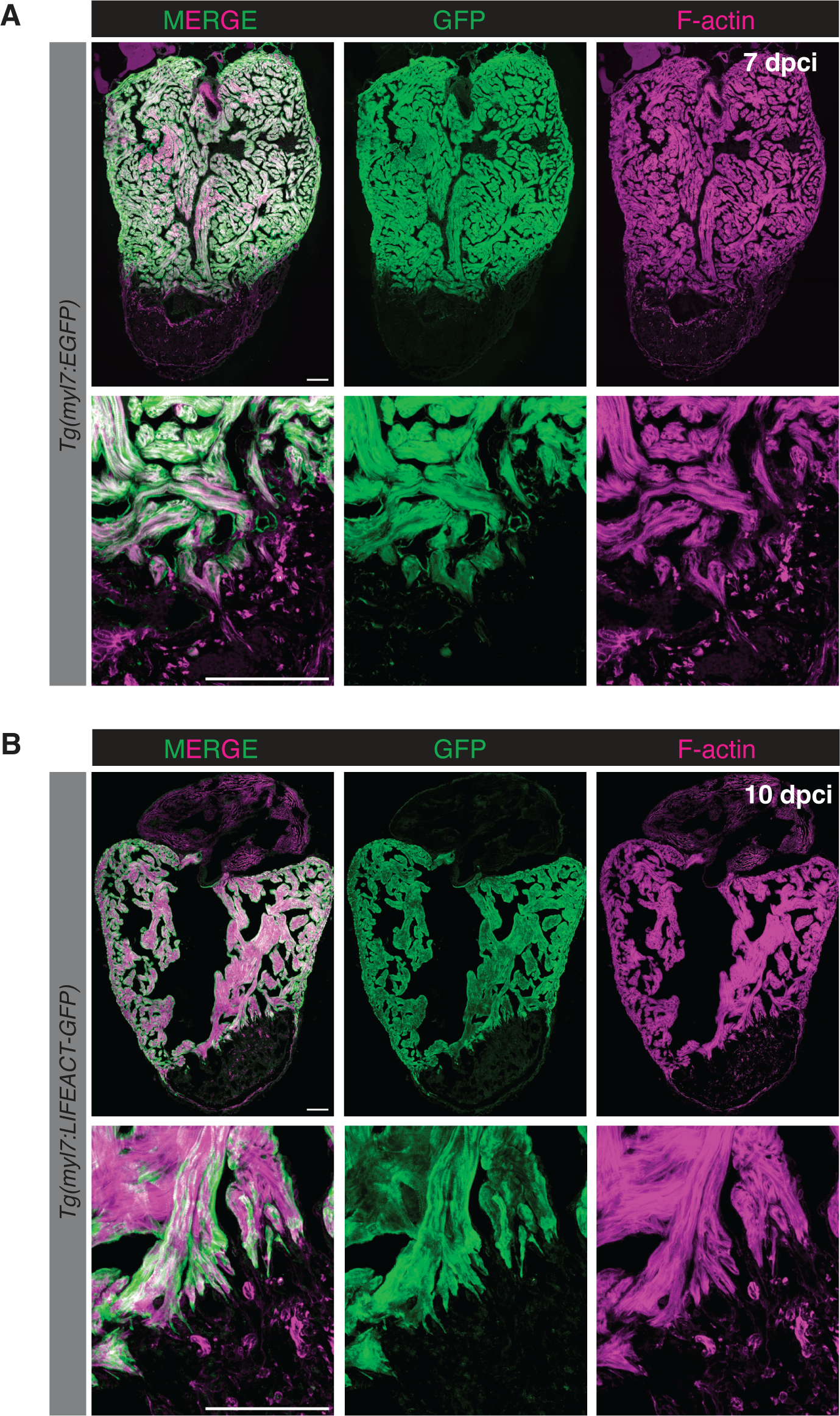
Phalloidin staining overlaps with CM-specific transgene expression. **(A)** Phalloidin and GFP immunostaining in *Tg(myl7:EGFP)* ventricles at 7 dpci. **(B)** Phalloidin and GFP immunostaining in *Tg(myl7:LIFEACT-GFP)* ventricles at 10 dpci. Scale bars: 100 μm.

**Supp. Figure 2:**
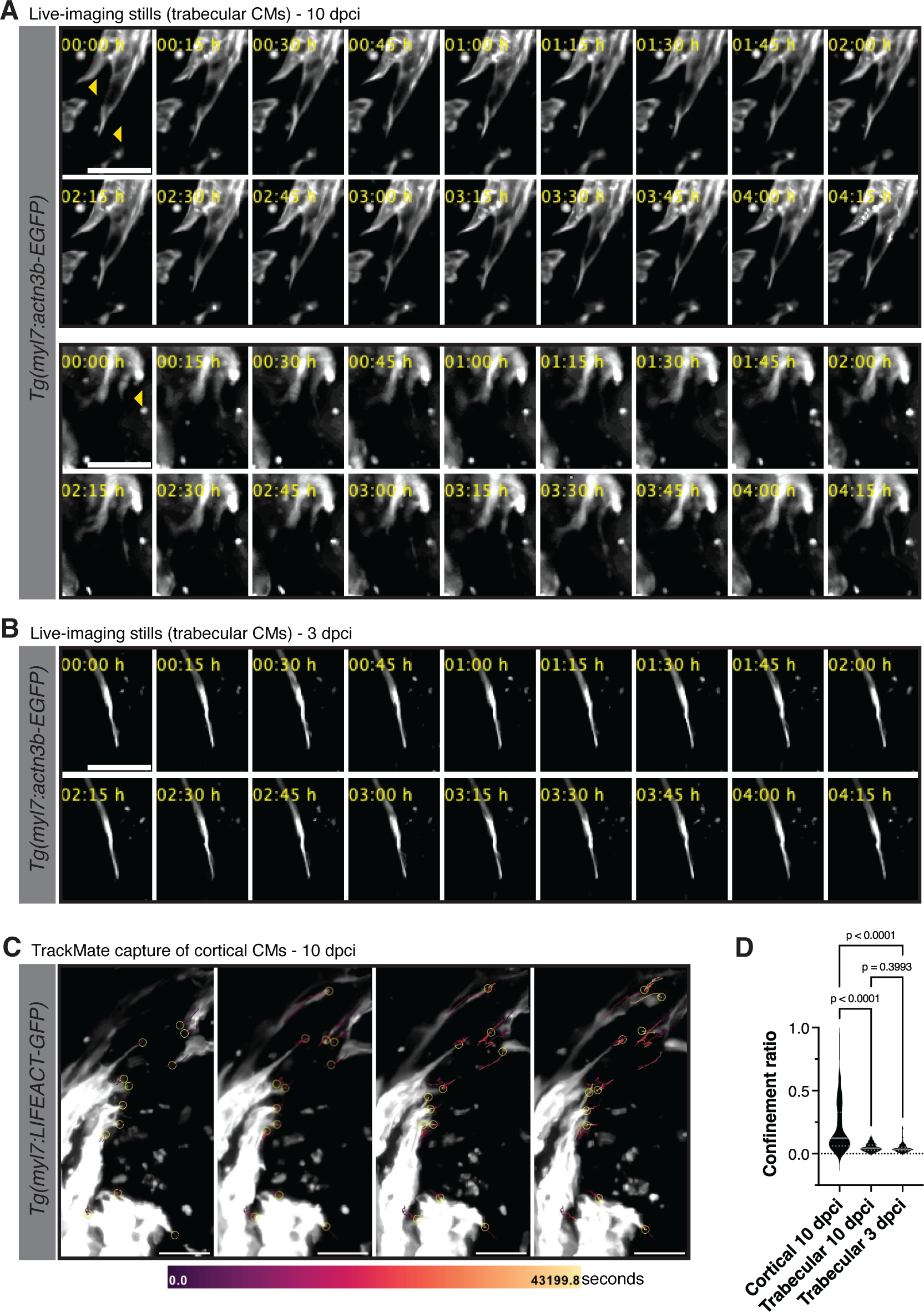
Time-lapse imaging and tracking of trabecular CM protrusions. **(A)** Time-lapse imaging of *Tg(myl7:actn3b-GFP)*+ trabecular CMs at 10 dpci. Yellow arrowheads point to CM protrusions that display a net positive migration into the injured area. **(B)** Time-lapse imaging of *Tg(myl7:actn3b-GFP)*+ trabecular CMs at 3 dpci. **(C)** TrackMate capture of *Tg(myl7:LIFEACT-GFP)*+ cortical CM protrusions at 10 dpci. Yellow circles denote the ends of CM protrusions that were tracked over a 12-hour time-lapse imaging period. Tracks were color-coded based on time. **(D)** Quantification of confinement ratio in CM protrusions at the border zone in *Tg(myl7:LIFEACT-GFP)*+ cortical CMs at 10 dpci and *Tg(myl7:actn3b-EGFP)*+ trabecular CMs at 3 and 10 dpci. For cortical CMs, n=114 CMs were tracked from 6 ventricles; for trabecular CMs at 10 dpci, n=84 CMs were tracked from 4 ventricles; for trabecular CMs at 3 dpci, n=75 CMs were tracked from 5 ventricles. P-values were calculated using a Kruskal-Wallis test with Dunnett’s multiple comparisons test. Scale bars: 20 μm.

**Supp. Figure 3:**
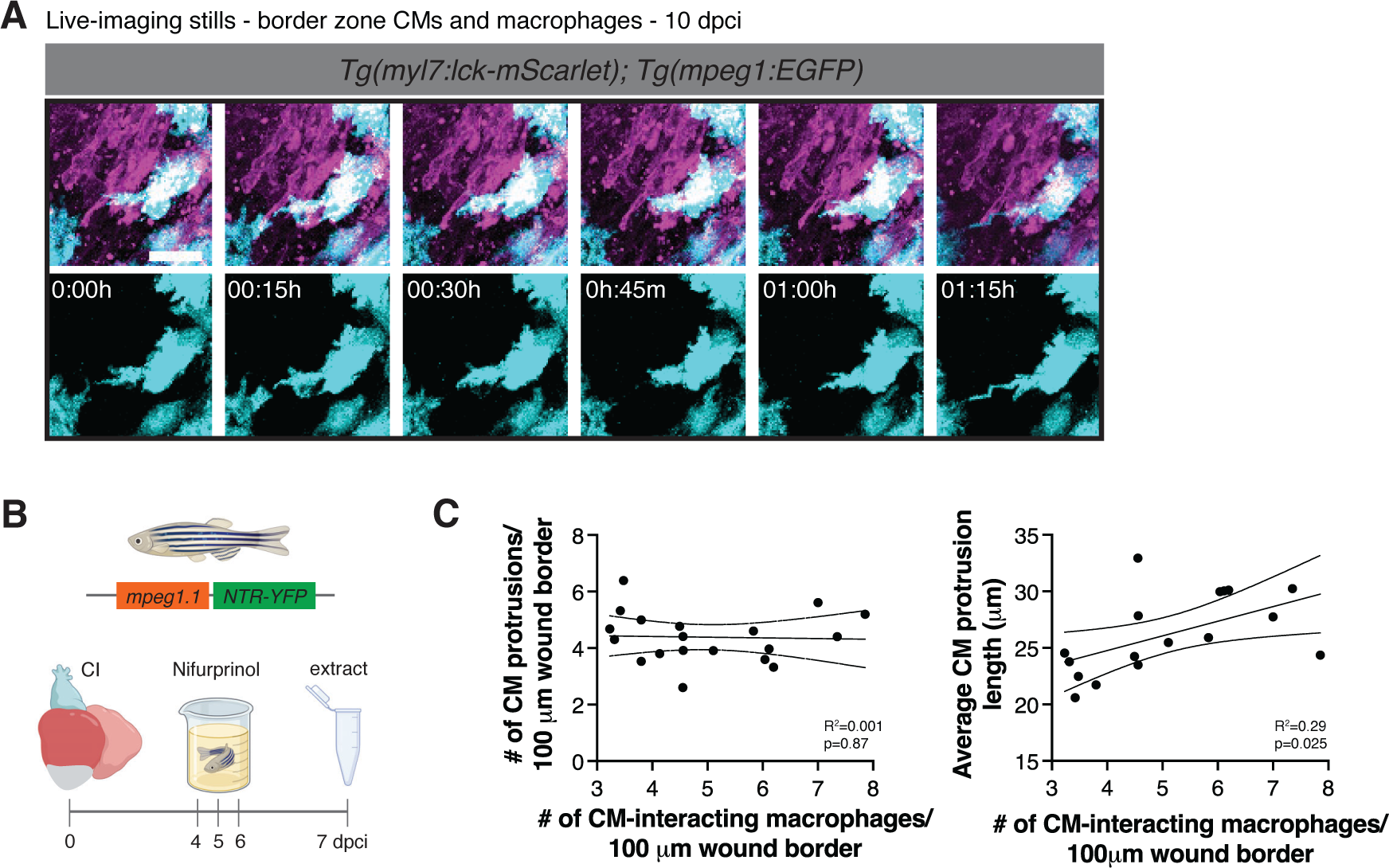
A decrease in macrophage presence affects CM protrusion in the regenerating zebrafish heart. **(A)** Time-lapse imaging of *Tg(myl7:lck-mScarlet); Tg(mpeg1:EGFP)* ventricular sections at the border zone at 7 dpci. **(B)** Schematic illustrating the macrophage ablation experimental set-up in *Tg(mpeg1.1:NTR-YFP)* ventricles. **(C)** Correlation between the number of CM-interacting macrophages and the number and length of CM protrusions at 7 dpci from n=11 DMSO-treated and n=8 nifurpirinol-treated *Tg(mpeg1:EGFP)* ventricles. Dashed lines indicate 95% confidence intervals of the best-fit line and p-values were calculated using simple linear regression. Scale bar: 10 μm.

**Supp. Figure 4:**
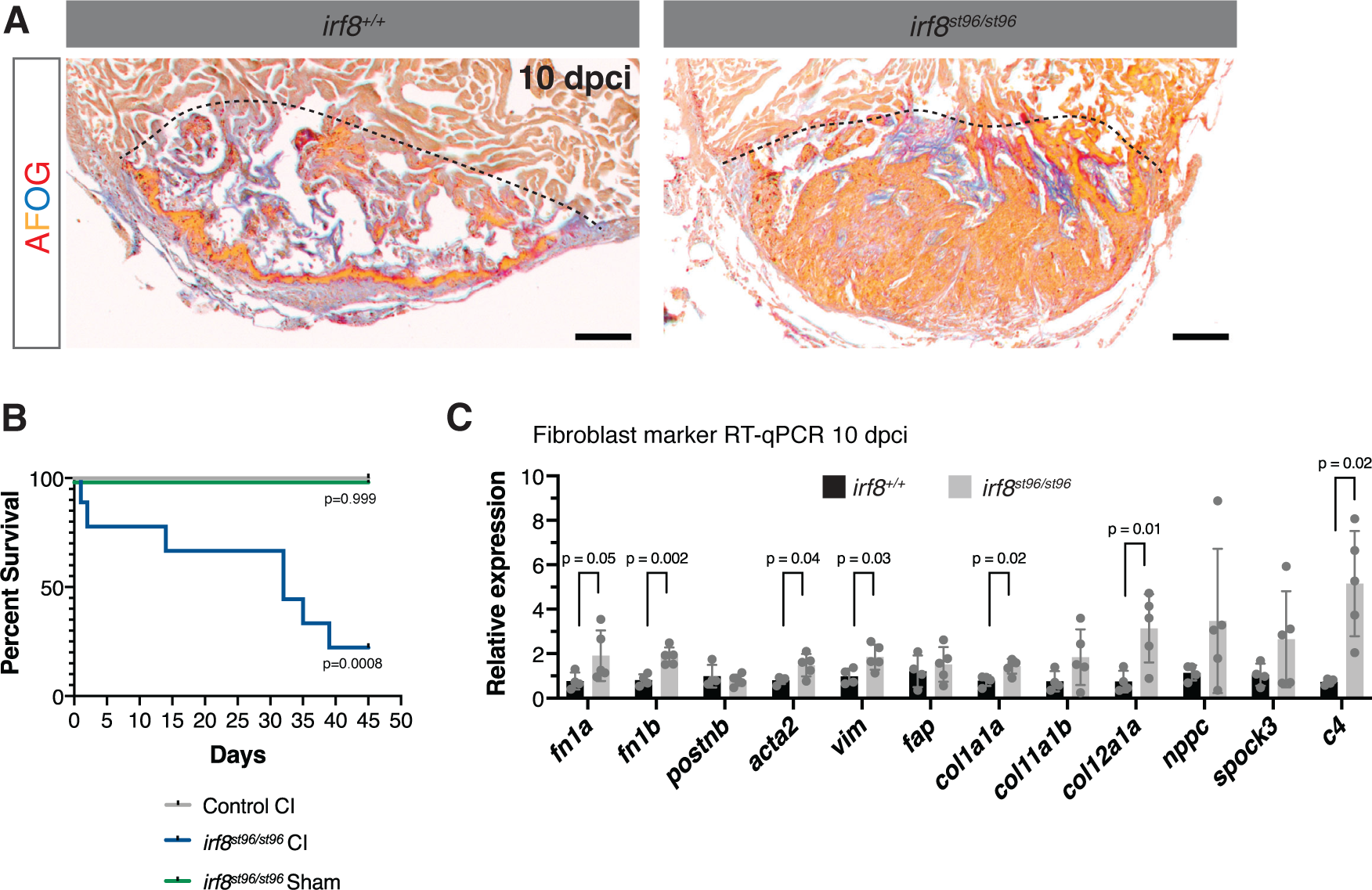
Absence of macrophages results in defects in scar composition, CM protrusion and organismal survival. **(A)** Acid fuchsin orange G (AFOG) staining in *irf8^st96/st96^* and wild-type sibling ventricles at 10 dpci. Black dashed lines indicate the approximate wound border. **(B)** Kaplan-Meier survival curve of wild-type control (n=9 fish) and *irf8^st96/st96^* mutant fish after sham (n=9 fish) or cryoinjury (n=9 fish). P-values compared to the control CI group were calculated using the Log-rank (Mantel-Cox) test. **(C)** RT-qPCR analysis of fibroblast marker genes in *irf8^st96/st96^* mutant (n=5) and wild-type sibling (n=4) ventricles at 10 dpci. P-values were calculated using an unpaired t-test or Mann-Whitney test (*c4*). Scale bars: 100 μm.

**Supp. Figure 5:**
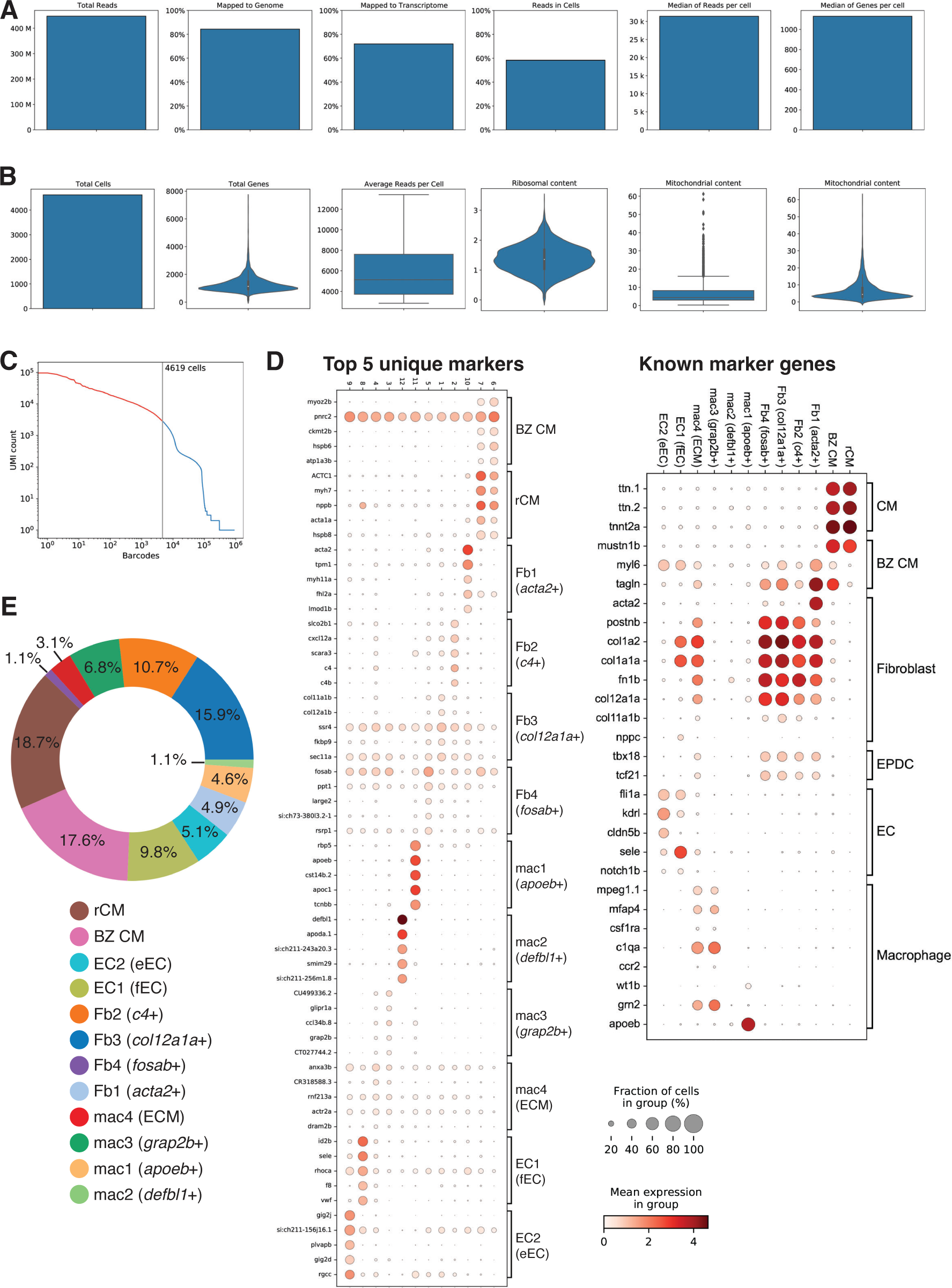
Quality control and initial analyses of scRNA-seq from border zone cells at 7 dpci. **(A) (B)** Quantitation and quality control metrics of scRNA-seq data, including total reads, mapped reads, median reads/genes per cell, ribosomal content, and mitochondrial content. **(C)** Knee plot of scRNA-seq data displaying the cut-off used for further analysis. **(D)** Dot plot of the top 5 unique marker genes for each cluster from the border zone scRNA-seq at 7 dpci (left). Dot plot of known marker genes from CMs, border zone CMs (BZ CMs), fibroblasts, epicardial-derived cells (EPDC), endothelial/endocardial cells (EC), and macrophages in the scRNA-seq data at 7 dpci (right). Red color denotes the mean expression within each group and size of the circle represents the fraction of cells expressing each gene within the group. **(E)** Pie chart illustrating the fraction of cells within each cluster as a percentage of all cells.

**Supp. Figure 6:**
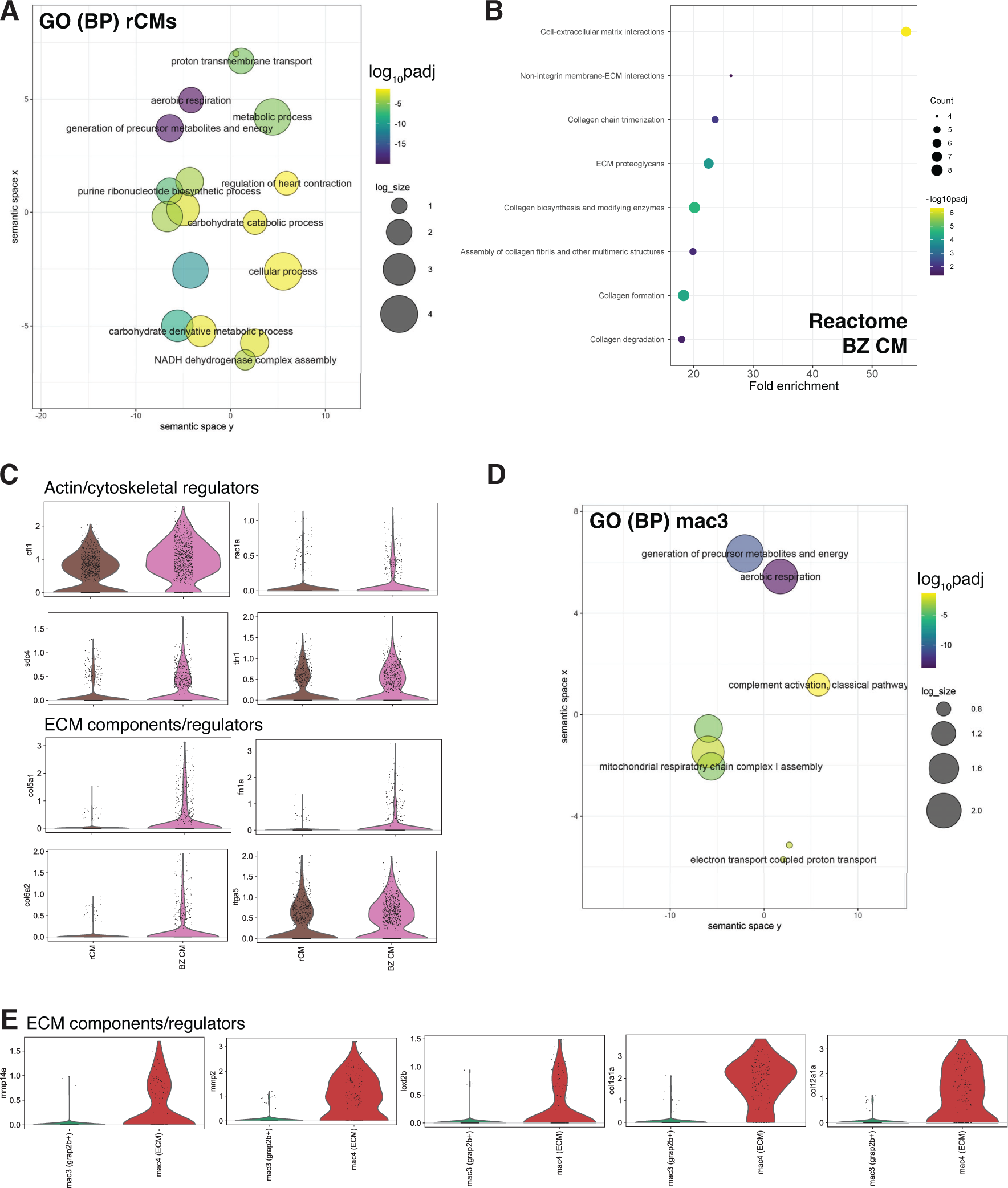
Gene ontology analysis of CMs and macrophages in border zone scRNA-seq at 7 dpci. **(A)** Gene ontology (GO) analysis of biological processes (BP) in differentially expressed genes enriched in remote CMs compared to border zone CMs. log_10_padj was calculated in comparison to all genes in the zebrafish genome and log_size corresponds to the log_10_(number of annotations for the GO Term ID in zebrafish from the EBI GOA Database). **(B)** Reactome pathway enrichment in genes upregulated in border zone CMs compared to remote CMs. Count refers to the number of genes included in each Reactome pathway term. **(C)** Violin plots of expression of genes involved in actin cytoskeleton and ECM regulation in rCMs vs. BZ CMs clusters from scRNA-seq at 7 dpci. **(D)** Gene ontology (GO) analysis of biological processes (BP) in differentially expressed genes enriched in mac3 compared to mac4(ECM). log_10_padj was calculated in comparison to all genes in the zebrafish genome and log_size corresponds to the log_10_(number of annotations for the GO Term ID in zebrafish from the EBI GOA Database). **(E)** Violin plots of expression of genes involved in actin cytoskeleton and ECM regulation in mac4 (ECM) vs. mac3 clusters from scRNA-seq at 7 dpci.

**Supp. Figure 7:**
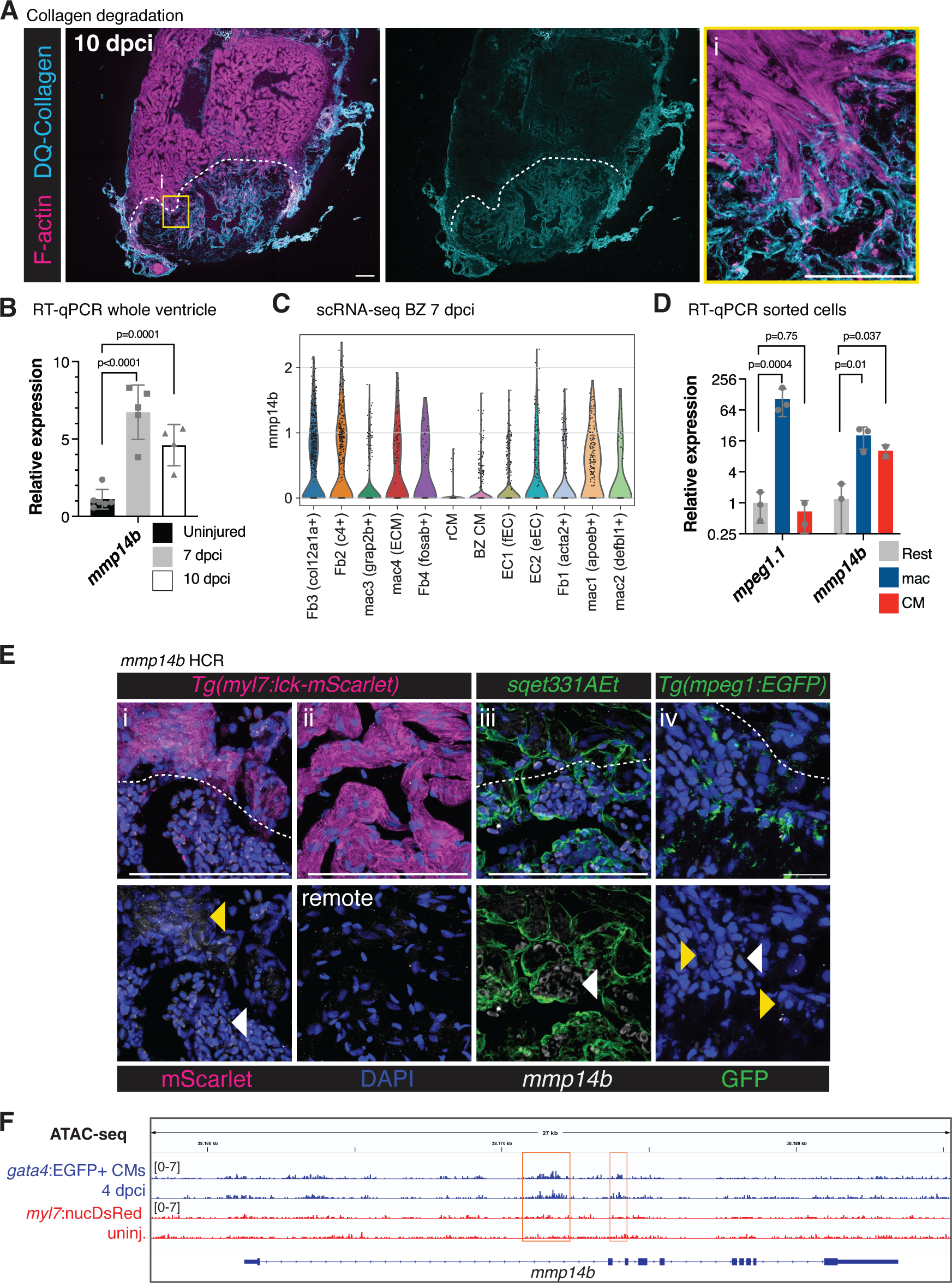
*mmp14b* expression in regenerating zebrafish hearts. **(A)** Incubation of fresh frozen sections with DQCollagen to fluorescently label collagenase activity in combination with phalloidin staining of F-actin in wild-type ventricles at 10 dpci. Yellow box denotes the zoomed image (i). White dashed lines indicate the approximate wound border. Images are representative of n=6 ventricles. **(B)** RT-qPCR analysis of *mmp14b* expression in whole uninjured ventricles (n=6) and in ventricles at 7 (n=5) and 10 dpci (n=4). P-values were calculated using one-way ANOVA and Dunnett’s multiple comparisons test. **(C)** Violin plots of *mmp14b* expression in cell clusters from the border zone scRNA-seq at 7 dpci. **(D)** RT-qPCR analysis of *mmp14b* expression in isolated CMs (n=2 biological replicates), macrophages (n=3 biological replicates), and non-CM/non-mac (n=3 biological replicates) sorted by fluorescent activated cell sorting (FACS) from *Tg(mpeg1:EGFP; myl7:mKATE-CAAX)* ventricles at 7 dpci. P-values were calculated using one-way ANOVA and Dunnett’s multiple comparisons test. **(E)** *mmp14b* expression in *Tg(myl7:lck-mScarlet), sqet331Aet,* and *Tg(mpeg1:EGFP)* ventricles, marking cardiomyocytes (i, ii), endocardial cells (iii), and macrophages (iv), respectively, at 10 dpci by *in situ* hybridization chain reaction (HCR). White dashed lines indicate the approximate wound border. Yellow arrowheads point to *mmp14b* HCR signal in border zone CMs and macrophages, and white arrowheads point to HCR signal in potential fibroblasts. **(F)** ATAC-seq tracks from regenerating *gata4*:EGFP+ CMs at 4 dpci and uninjured *myl7*:nucDsRed+ CMs showing CM-specific chromatin accessibility at the *mmp14b* locus. Scale bars: 100 μm and 20μm in **(E, iv)**.

**Supp. Figure 8:**
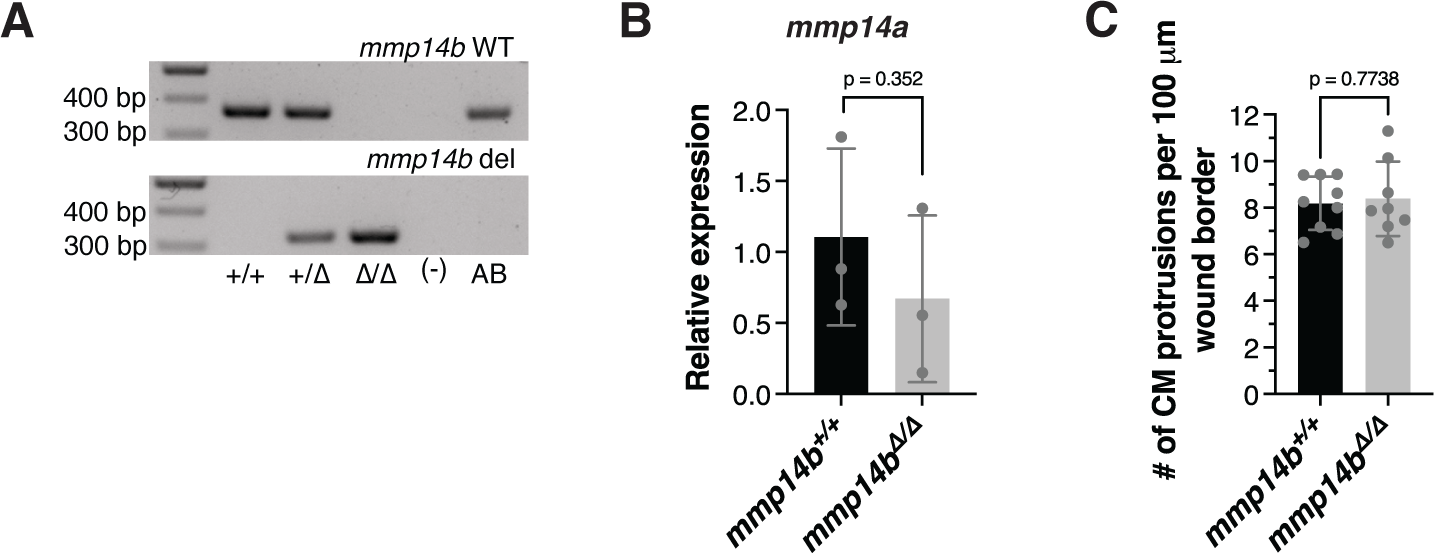
*mmp14b* mutant embryos display no genetic compensation from *mmp14a*. **(A)** Genotyping of genomic DNA from *mmp14b^+/+^*, *mmp14b^+/11^*, and *mmp14b^11/11^* fish with primers specific for the *mmp14b* wild-type allele and *mmp14b* full locus deletion allele. **(B)** RT-qPCR analysis of *mmp14a* expression from pools of *mmp14b^+/+^* (n=3 pools from 3 separate clutches of embryos) and *mmp14b^11/11^* (n=3 pools from 3 separate clutches of embryos) embryos at 48 hpf. **(C)** Quantification of the number of CM protrusions in *mmp14b^11/11^* mutant (n=8) and wild-type sibling (n=9) ventricles at 10 dpci. P-values were calculated using an unpaired t-test.

**Supp. Figure 9:**
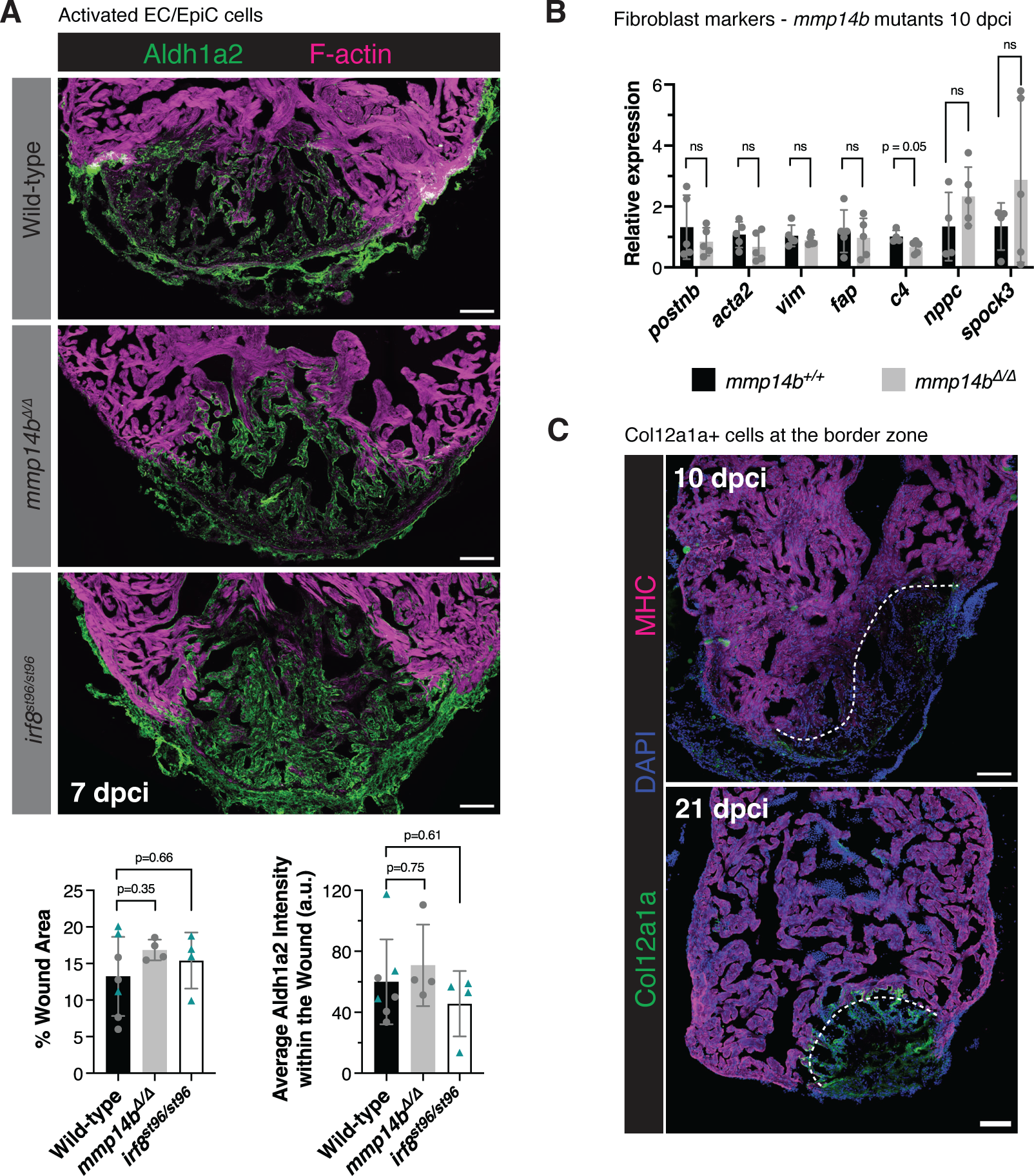
Endocardial and fibroblast response in *mmp14b* mutants. **(A)** Phalloidin and Aldh1a2 immunostaining in wild-type, *mmp14b^11/11^* mutant, and *irf8^st96/st96^* mutant ventricles at 7 dpci. Quantification of the percent Aldh1a2+ staining within the wound area and average Aldh1a2 intensity within the wound in wild-type (n=7), *mmp14b^11/11^* mutant (n=4), and *irf8^st96/st96^* mutant (n=4) ventricles at 7 dpci are shown in the lower panel. P-values were calculated using ordinary one-way ANOVA and Dunnett’s multiple comparisons test. Wild-type siblings from *mmp14b* and *irf8* mutants were grouped together. **(B)** RT-qPCR analysis of fibroblast gene expression in *mmp14b^11/11^* mutant (n=5) and wild-type sibling (n=5) ventricles at 10 dpci. ns, not significant. P-values were calculated using an unpaired t-test or Mann Whitney test (*spock3*). **(C)** Col12a1a and Myosin heavy chain (MHC) immunostaining in ventricles at 10 and 21 dpci. White dashed lines indicate the approximate wound border. Scale bars: 100 μm.

**Supplementary Video 1: Time-lapse imaging of cortically-located (*Tg(myl7:LIFEACT-GFP)*+) CM protrusion into the injured area at 10 dpci.**

**Supplementary Video 2: Tracking of cortically-located CM protrusions at 10 dpci using TrackMate.**

**Supplementary Video 3: Time-lapse imaging of trabecular (*Tg(myl7:actn3b-EGFP)*+) CM protrusion into the injured area at 10 dpci.**

**Supplementary Video 4: Time-lapse imaging of trabecular (*Tg(myl7:actn3b-EGFP)*+) CM protrusion into the injured area at 3 dpci.**

**Supplementary Video 5: Time-lapse imaging of border zone *Tg(mpeg1:EGFP)*+ cells at 10 dpci.**

**Supplementary Video 6: Time-lapse imaging of *Tg(mpeg1:EGFP)*+ cells in the injury at 10 dpci.**

